# Therapeutic blockade of Activin-A improves NK cell function and anti-tumor immunity

**DOI:** 10.1101/454801

**Authors:** Jai Rautela, Laura F. Dagley, Iona S. Schuster, Soroor Hediyeh-Zadeh, Rebecca B. Delconte, Joseph Cursons, Robert Hennessy, Dana S. Hutchinson, Craig Harrison, Carolina C. de Oliveira, Eric Vivier, Andrew I. Webb, Mariapia A. Degli-Esposti, Melissa J. Davis, Nicholas D. Huntington, Fernando Souza-Fonseca-Guimaraes

## Abstract

Natural killer (NK) cells are innate lymphocytes that play a major role in immunosurveillance against tumor initiation and metastasis spread. Signals and checkpoints that regulate NK cell fitness and function in the tumor microenvironment are not well defined. Transforming grow factor (TGF)-β is a recognized suppressor of NK cells that inhibits IL-15 dependent signaling events and induces cellular transdifferentiation, however the role of other SMAD signaling pathways in NK cells is unknown. In this report, we show that NK cells express the type I Activin receptor, ALK4, which upon binding its ligand Activin-A, phosphorylates SMAD2/3 to efficiently suppress IL-15-mediated NK cell metabolism. Activin-A impairs human and mouse NK cell proliferation and downregulates intracellular granzyme B levels to impair tumor killing. Similar to TGF-β, Activin-A also induced SMAD2/3 phosphorylation and drove NK cells to upregulate several ILC1-like surface markers including CD69, TRAIL and CD49a. Activin-A also induced these changes on TGF-β receptor deficient NK cells, highlighting that Activin-A and TGF-β are independent pathways that drive SMAD2/3-mediated NK cell suppression. Finally, therapeutic inhibition of Activin-A by Follistatin significantly slowed orthotopic melanoma growth in mice. These data highlight independent SMAD2/3 pathways target NK cell fitness and function and identify a novel therapeutic axis to promote tumor immunity.

**One Sentence Summary:** Activin-A can directly inhibit NK cell effector functions, promote NK cells transdifferentiation into ILC1-like cells and suppress anti-melanoma immunity.

## Introduction

NK cells are implicated in controlling tumor initiation and metastasis, and as such these effector functions are being utilized in cancer clinical trials (Krasnova et al., 2017; Souza-Fonseca-Guimaraes, 2016). Several biological factors have been reported to directly suppress or limit NK cell function via a range of mechanisms in malignant diseases. These include tumor derived metabolites such as adenosine (Beavis et al., 2013a; Beavis et al., 2013b) or enzymes such as indoleamine 2,3-dioxygenase-1 (IDO1 or IDO) (Liu et al., 2010), upregulation of cytokine-inducible SH2-containing protein (CIS, encoded by *Cish*) (Delconte et al., 2016a), downregulation of anti-apoptotic Bcl2 and Mcl1 proteins (Huntington et al., 2007; Sathe et al., 2014; Viant et al., 2017), modulation of NKG2D expression (Raulet et al., 2013), expression of inhibitory receptors for MHC-I (Benson and Caligiuri, 2014; Rahim and Makrigiannis, 2015), and transforming growth factor beta (TGF-β) (Gao et al., 2017; Viel et al., 2016).

TGF-β is a secreted protein that has three different isoforms, TGF-β1, TGF-β2 and TGF-β3. These isoforms are expressed in different tissues, with TGF-β1 being the most abundant isoform and a highly potent modulator of the immune system (Kubiczkova et al., 2012). TGF-β1 is a pleiotropic cytokine produced by several cell subtypes (including tumor cells, regulatory T cells, stromal fibroblasts, and various myeloid cell subsets) and is an important suppressor of NK cell anti-tumor and anti-metastatic functions in experimental tumor models (Ghiringhelli et al., 2005; Souza-Fonseca-Guimaraes and Smyth, 2013; Viel et al., 2016). TGF-β1 reduces NK cell priming and activation, including both cytotoxicity and cytokine production, by suppressing the mTOR pathway (Bellone et al., 1995; Viel et al., 2016). In line with this, a previous study demonstrated that anti-TGF-β neutralizing antibodies increase NK cell effector functions *in vitro* and *in vivo* (Arteaga et al., 1993).

TGF-βRI and II are transmembrane receptors associated with serine/threonine kinases that, upon ligand binding, mediate the phosphorylation of the SMAD family of transcription factors (Shi and Massague, 2003). Although TGF-β is the best described inducer of this pathway, other factors (Activin A, Activin B, myostatin (GFF8), GDF11, Nodal and bone morphogenetic proteins (BMPs)), which play a critical role in bone and muscular development) can also trigger the phosphorylation of SMAD2/3 (Lin et al., 2016; Loomans and Andl, 2016; Pasteuning-Vuhman et al., 2017). Activin-A binds to a set of receptors (ALK4, ALK7, ActRIIA, and ActRIIB) distinct from those used by the TGF-β isoforms (Loomans and Andl, 2016). Activin-A and its receptor dimer comprising ALK4 and ACVR2A/B play a critical role in muscular development (Pasteuning-Vuhman et al., 2017), and can be produced in large amounts by dendritic cells (DCs) or following acute inflammation (Jones et al., 2004; Robson et al., 2008).

In immune cells, while TGF-β signaling is a critical step in CD4^+^ T cell differentiation into CD4^+^ Foxp3^+^ regulatory cells (Tregs), Activin-A has also been demonstrated to induce Foxp3 expression and promote Treg conversion (Huber et al., 2009; Semitekolou et al., 2009). In breast and ovarian cancer cells, Activin-A signaling was previously shown to induce epithelial-mesenchymal transition (EMT), a malignant cellular reprograming that is characteristic of TGF-β signaling (Bashir et al., 2015; Basu et al., 2015). In addition, Activin signaling genes were previously shown to be upregulated in breast cancer cell EMT (Cursons et al., 2015). High circulating Activin-A levels were also shown to be associated with tumor progression and poor prognosis in lung cancer patients (Hoda et al., 2016). More recently, Activin signaling was shown to reprogram macrophages into protumorigenic subsets which promote skin carcinogenesis (Antsiferova et al., 2017). The biological activity of Activin-A is negatively regulated by an endogenous inhibitor known as Follistatin (FST) and also at the cell surface by *TGFBR3* (betaglycan) (Walton et al., 2012). Recently, studies have shown a correlation between reduced FST expression and lower survival rates among breast cancer and cutaneous melanoma patients (Donovan et al., 2017; Seachrist et al., 2017). Similarly, low expression of *TGFBR3* was found to be prognostic for poor survival in renal cell carcinoma (Nishida et al., 2018). Previous work has also shown that dendritic cell (DC)-derived Activin-A reduced cytokine production and cellular activation of peripheral blood human NK cells (Robson et al., 2009). Although previous results have hinted at the involvement of alternate SMAD2,3 pathways in regulating human NK cell function, the precise mechanism and influence on NK cell biology is still to be elucidated.

The broader impact of TGF-β signaling on group 1 innate lymphoid cells (consisting of conventional NK cells (cNK) and type 1 innate lymphoid cells (ILC1) (Spits et al., 2013)), was recently demonstrated in the salivary gland and has renewed interest in the precise effect of TGF-β on NK cells function within the tumor microenvironment (Cortez et al., 2016). We recently identified a novel unidirectional reprogramming of cNK cells into ILC1-like cells driven by TGF-β in the tumor microenvironment. Transgenic mouse models with ablated or constitutive TGF-β signaling specifically in NKp46^+^ cells had reduced numbers of ILC1s and cNK cells, respectively. TGF-β signaling was also capable of driving cNK cells to acquire a transitional ILC1-like phenotype both *in vitro* and *in vivo* by downregulating Eomesodermin (Eomes), and upregulating tissue residency-related markers such as tumor necrosis factor-related apoptosis inducing ligand (TRAIL), and the collagen binding protein CD49a in the tumor microenvironment (Gao et al., 2017). Intriguingly, a population of NKp46^+^CD49a^+^ cells was still observed in the tumor microenvironment of NKp46^cre/+^TgfbRII^fl/fl^ mice, suggesting a minor TGF-β-independent pathway for ILC1 differentiation may exist. Here we describe for the first time that Activin-A acts as an alternate SMAD signaling pathway to suppress NK cell metabolism and cytotoxicity, ultimately inducing tissue residency features on NK cells.

## Results

### Functional Activin-A receptor expression on NK cells

Receptors for Activin-A are classified as type 1 (ACVR1B or ALK4, and ACVR1C or ALK7) and type 2 (ACVR2A, and ACVR2B) receptors (Lotinun et al., 2012). Human NK cells are reported to express mRNA for Activin-A receptors (ALK4, ACVRIIA and ACVRIIB) and potentially respond to Activin-A (Robson et al., 2009). Using available RNAseq data (Delconte et al., 2016b; Holmes et al., 2014; Seillet et al., 2014), we compared the mRNA levels of Activin receptors in different murine NK cell subsets: progenitors (pre-pro NK cells, and NKP cells), immature (CD27^+^CD11b^neg^) NK cells, mature (CD27^neg^CD11b^+^) NK cells, and liver tissue resident or ILC1 (TRAIL^+^CD49b^neg^) cells. Notably, all subsets express ALK4 mRNA which declines as NK cells mature – paralleling the expression pattern of TGF-βRII (fig. 1A) and consistent with our previous report (Viel et al., 2016). However, we failed to detect a clear mRNA signal for receptors ACVR2A, ACVR2B and ACVR1C receptors in these NK cell subsets (fig. S1).

**Figure 1.**
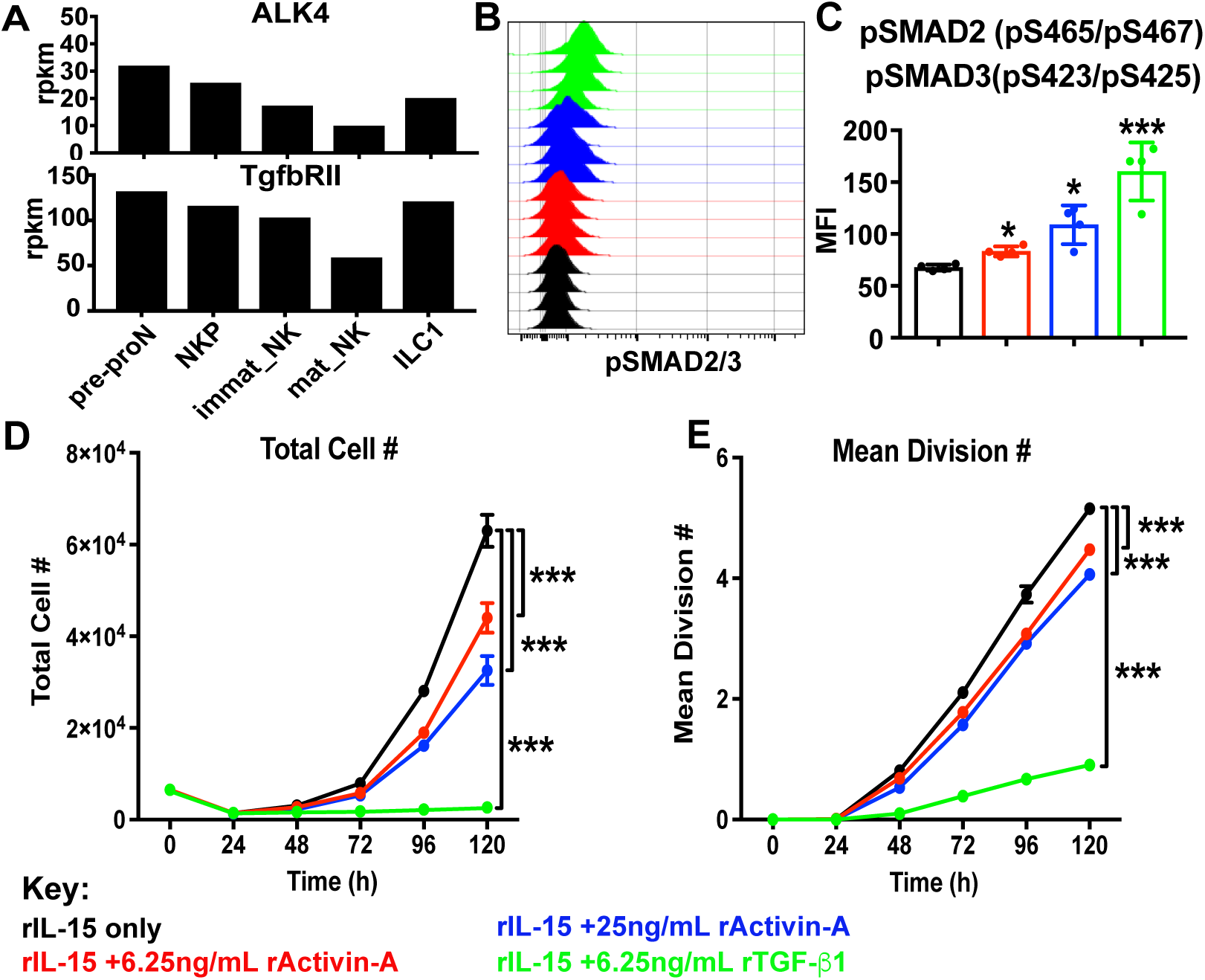
Activin receptor expression and responsiveness in NK cells. (A) mRNA abundance of ALK4 and TGF-βRII (TgfbRII) in pre-pro-NK cells, NK cell progenitors (NKP), immature (immat_NK), mature (mat_NK) cells, and liver ILC1s, (B-C) Purified NK cells were challenged with the indicated concentrations of rActivin-A or TGF-β1 for 1 h, and pSMAD2/3 was stained for FACS. (B) Overlay represents individual biological replicates. (C) MFI comparison of pSMAD2/3 stain. Results are expressed as the mean + SEM, and unpaired T test was used for comparative statistical analysis, where *p < 0.05, ***p < 0.001. (F-G) CTV-labelled NK cells were cultured in presence of rIL-15 in addition to rTGF-β1 or rActivin-A as indicated. For each condition at 24 h interval, the total live cells were enumerated and population average division number was calculated. Results are representative from one representative biological replicate in triplicate from three independent experiments, and two-way ANOVA used for statistical analysis where ***p < 0.001.

A challenge in the field of TGF-β research is the highly conserved homology of this cytokine between species (Herpin et al., 2004). Due to the presence of bovine TGF-β in conventional media containing fetal calf serum (FCS) (fig. S2), our *in vitro* assays used media free from animal-derived components to compare the effects of recombinant TGF-β versus Activin-A. Using recombinant Activin-A (rActivin-A), we observed that purified mouse NK cells phosphorylated the SMAD2/3 complex in a dose-dependent manner, but not to the same degree as recombinant TGF-β1 (rTGF-β1) (fig. 1B-C). Similarly, both rTGF-β1 and rActivin-A (particularly the highest concentration tested) inhibited Ki67 expression (data not shown) and NK cell proliferation in response to IL-15 (fig. 1D-E). We found that neither ligand induced NK cell death (as shown by total cohort number, fig. S3) (Viant et al., 2017), but rather suppressed the rate at which NK cells divide (fig 1F-G). Strikingly, rTGF-β1 concentrations as low as 0.1 ng/mL were enough to suppress NK cell proliferation (fig. S4) more efficiently than 1 µM of 5’-(N-Ethylcarboxamido)adenosine (NECA), a well described inhibitor of NK cell functions (Beavis et al., 2013a; Young et al., 2017). These results reveal that NK cells respond to Activin-A, albeit with lower SMAD2/3 phosphorylation and a less striking impact on proliferation compared to TGF-β.

### Activin signaling drives tissue residency features on NK cells

Lymphocytic tissue residency is characterized by decreased proliferation and expression of tissue residence markers such as the collagen binding receptor CD49a (or ITGA1, or VLA-1) which functions as a cellular anchor (Bank et al., 1994). *In vivo* parabiosis experiments have demonstrated that tissue resident Eomes^neg^CD49a^+^ NK cells, or ILC1s, are unable to migrate between parabiont mice, while conventional Eomes^+^CD49a^neg^ NK cells (cNK) migrate freely (Sojka et al., 2014). Recently, we revealed that rTGF-β1 simultaneously upregulates CD49a expression while downregulating Eomes expression in cNK cells *in vitro*. In addition, although TGF-βR-deficient NKp46^+^ cells are completely unresponsive to rTGF-β1 *in vitro*, there is still clear CD49a expression on a subset of these cells *in vivo* (Gao et al., 2017). To investigate if cNK transdifferentiation (Eomes downregulation and CD49a upregulation) could be induced by rActivin-A similarly to rTGF-β1, we cultured highly purified (99-100% purity) splenic cNK cells (viable, CD3/CD19/F4-80/Ly6G/TCRβ^neg^, CD49a^neg^, NKp46^+^, NK1.1^+^, CD49b^+^) from NKp46^cre/+^TgfbRII^fl/fl^ mice, and corresponding littermate controls (NKp46^+/+^TgfbRII^fl/fl^), in animal component-free media supplemented with rActivin-A or rTGF-β1. Surprisingly, rActivin-A was as efficacious as rTGF-β1 at inducing the expression of CD49a expression on WT NK cells, and unlike TGF-β, also acted upon TgfbRII-deficient NK cells (fig. 2A&E). But in contrast, Activin-A signaling did not efficiently downregulate Eomes expression in murine NK cells, whereas rTGF-β1 did (fig. 2B&F).

**Figure 2.**
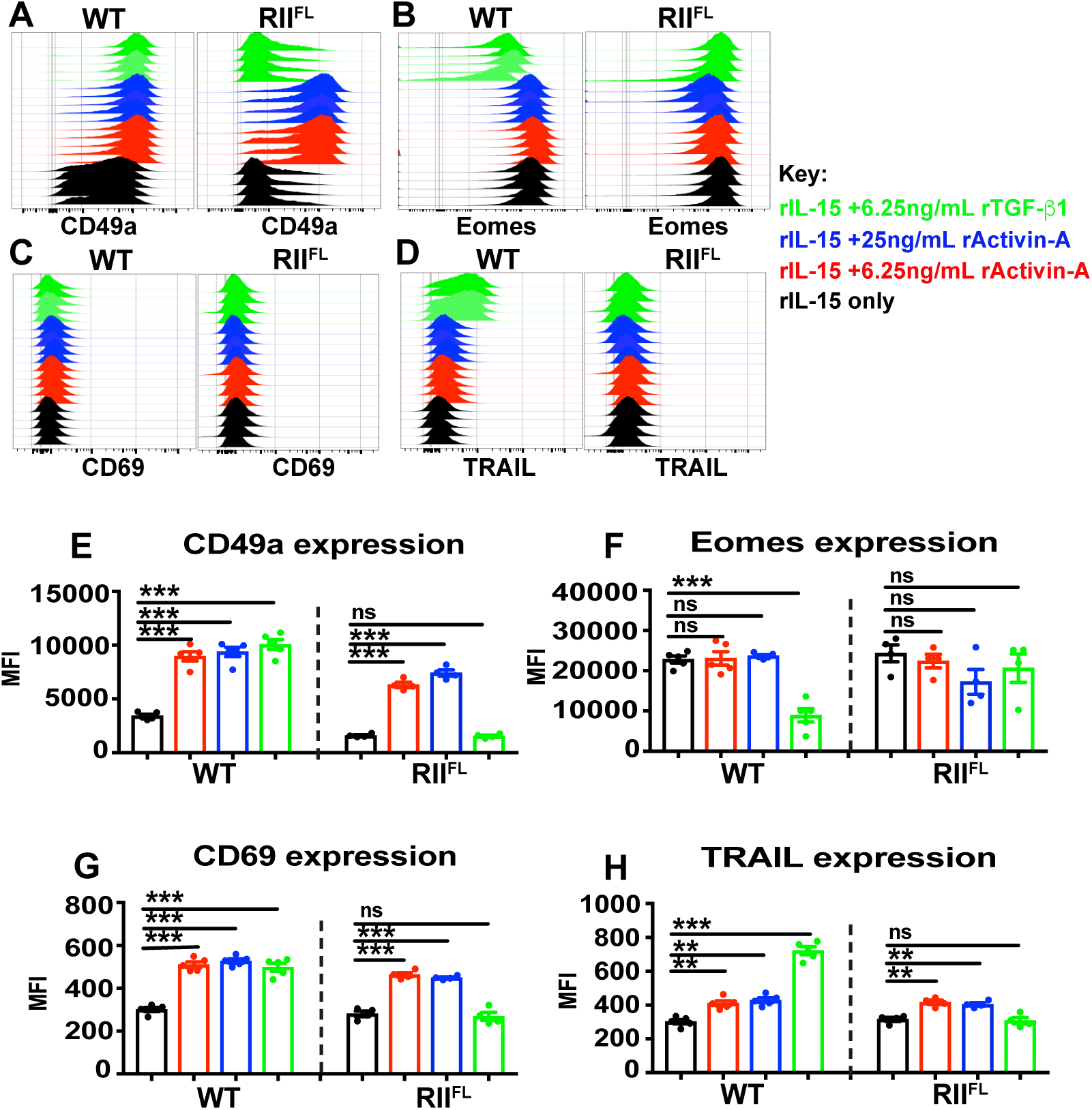
Activin-A induce transdifferentiation features in conventional NK cells. (A-H) Purified splenic cNK cells from spleens RII^FL^ (NKp46^cre/+^TgfbRII^fl/fl^) or respectively littermate control mice were cultured in 50 ng/mL rIL-15, with or without the indicated concentrations of rActivin-A or rTGF-β1 for 48 h. Surface CD49a (A and E), intracellular Eomes (B and F), surface CD69 (C and G) and surface TRAIL (D and H) expression were determined by flow cytometry. Each overlay represents one independent biological replicate of the indicated cytokine stimulation condition (A, B, C and D for CD49a, Eomes, CD69 and TRAIL respectively). Results are expressed as the mean ± SEM of MFI, each data point represents one independent biological replicate and one-way ANOVA was used for comparative statistical analysis, where *p < 0.05, **p < 0.01 and ***p < 0.001.

ILC1s or tissue resident NK cells, are not only defined by their expression of CD49a and Eomes-independent development but are also identified by their expression of the early activation marker CD69 and TNF-related apoptosis-inducing ligand (TRAIL) at steady state (Sojka et al., 2014). CD69-induction by TGF-β signaling and its correlation with tissue residency, is a well described phenomenon for T cell residency (Mackay et al., 2015a; Mackay et al., 2013; Mackay et al., 2015b). We observed that rActivin-A induced CD69 upregulation similarly to and independently of rTGF-β1 in cNK cells *in vitro* (fig. 2C&G). We previously showed that rTGF-β1 efficiently upregulates TRAIL in cNK cells *in vitro* (Gao et al., 2017), and TRAIL is a well-documented marker to defining liver ILC1s from fetal liver throughout ontogeny (Takeda et al., 2005). rActivin-A signaling was sufficient to upregulate TRAIL, although in this case, TRAIL expression was less efficiently induced by rActivin-A than rTGF-β1 (fig. 2D&H). These findings suggest that Activin signaling induces changes in NK cell phenotype reminiscent of those involved in NK cell transdifferentiation mediated by TGF-β.

### Activin signaling regulates NK cell activation and metabolism

Alteration in cellular morphology is a notable consequence of TGF-β signaling in NK cells, and is directly linked to inhibition of metabolism mediated by the molecular target of rapamycin (mTOR) (Viel et al., 2016). We used scanning electron microscopy to detect morphological characteristics with nanometer resolution, both with or without overnight rTGF-β1 incubation and interaction with MHC-I-deficient melanoma cells. As anticipated, we observed that TGF-β signaling impacted NK cell morphology and appeared to reduce NK cell adherence to their targets (fig. S5).

To gain insight into the pathways modulated by SMAD2/3 signaling in NK cells, we performed a global proteomics and transcriptomics analyses. (fig. 3, fig. 4A). To identify the mediators and effectors of TGF-β-induced SMAD2,3 signaling, we stimulated splenic NK cells from WT and TGF-βRII^FL^ mice (as background control) and performed RNAseq. Together with our previous data derived from intratumoral intILC1s, we could further refine the lists of 189 upregulated (fig. 3A) and 228 downregulated genes (fig. 3B), induced by TGF-β signaling. Among these lists of differentially-expressed genes, there was an abundance of cell cycle-related genes that were downregulated after TGF-β treatment (fig. 3C), further corroborating our findings in fig 1F-G. Global proteomic analysis of TGF-β treated WT NK cells revealed a reduction in the expression of proteins involved in cellular metabolism and cellular activation (fig 4A-B). Namely, *Tnfrsf9* (CD137), *Gzmb*, *Hmgcs1*, *Acly* and *Prf1* are all significantly downregulated in WT samples upon TGF-β1 stimulation, while their protein expression remains unchanged in the RII^FL^ samples. This list of downregulated proteins included (Also summarized in table S1): ATP citrate lyase (ACLY), which is critical in converting citrate to acetyl-CoA and links the metabolism of carbohydrates (Sun et al., 2010); hydroxymethylglutaryl-CoA synthase (HMGCS1), which condenses acetyl-CoA for further metabolic changes (Mathews et al., 2014); cytotoxic proteins as Perforin (*Prf1*) and Granzyme B (*Gzmb*), in agreement with our previous report (Viel et al., 2016) and the tumor necrosis factor receptor superfamily 9 (TNFRSF9, 4-1BB or CD137), a lymphocyte activation receptor known to enhance NK cell function (Kohrt et al., 2011; Kohrt et al., 2012; Rajasekaran et al., 2013).

**Figure 3.**
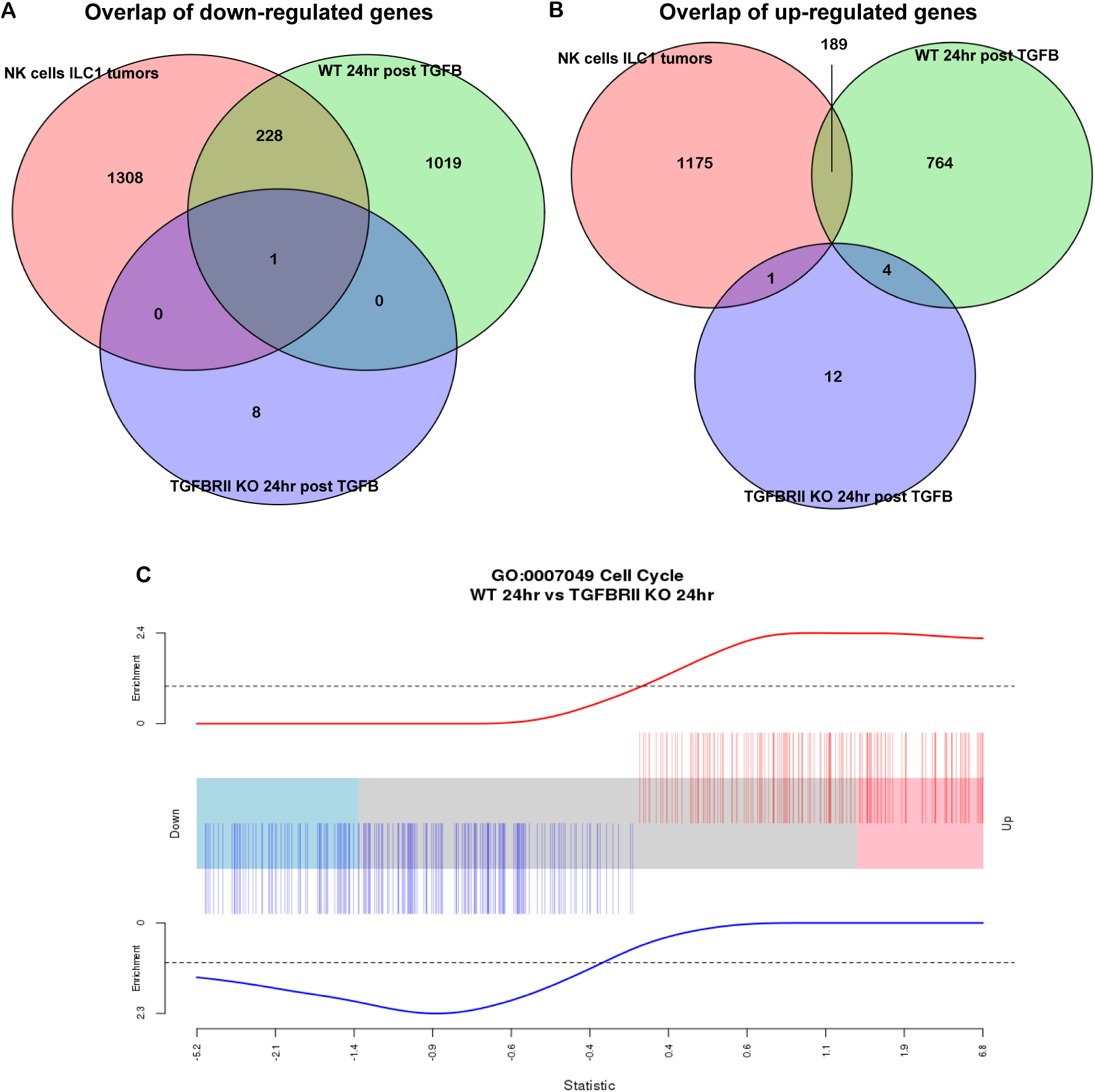
TGF-β signaling regulates cell cycle transcriptomics of NK cells. (A-B) Changes in the expression of cell cycle genes in WT cells versus TGF-βRII-deficient NK cells 24 h post rTGF-β1 treatment. Venn diagrams display the overlap between genes down (A) or up-regulated (B) in WT NK cells 24 h after rTGF-β1 treatment compared to unstimulated state *in vitro* with genes changing in the same direction as *in vivo* ILC1-like cells was determined. A signature was derived by excluding genes up or down-regulated in TFGBRII-KO NK cells 24 h after rTGF-β1 treatment. The obtained signature represents the genes that are most likely to be regulated by TGF-β signaling in NK cells both *in vitro* and *in vivo*. (C) Enrichment of the obtained refined gene sets was compared for significant changes in the “Cell Cycle” pathway analysis post 24 h rTGF-β1 stimulation.

**Figure. 4.**
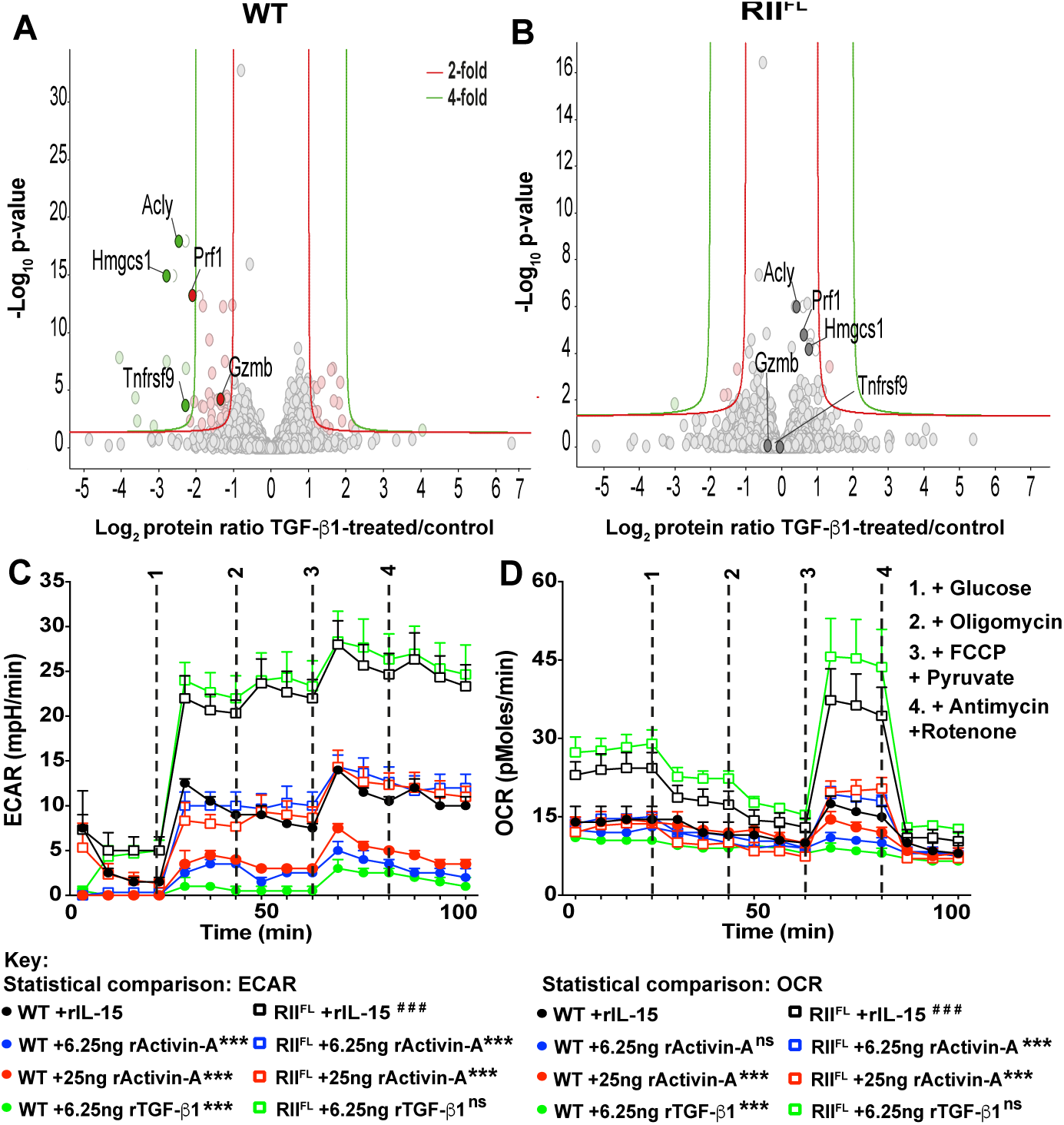
Activin-A signaling impacts in NK cell metabolism. (A-B) IL-15-expanded NK cells of WT or RII^FL^ mice were cultured for 24 h in presence of rIL-15 with or without rTGF-β1, and then processed for shotgun proteomics. Volcano plots illustrating the log_2_ protein ratios of proteins in cultured cells with and without TGF-β1 stimulation were compared for WT samples (A) and in RII^FL^ plus TGF-β1 stimulation relative to unstimulated RII^FL^ samples (B) following quantitative pipeline analysis. Proteins were deemed differentially regulated if the log_2_ fold change in protein expression was greater than 2-fold (red) or 4-fold (green) and a −log_10_ *p* value ≥ 1.3, equivalent to a *p* value ≤ 0.05. (C-D) IL-15-expanded NK cells were cultured overnight (20h) in presence of rIL-15 with or without rActivin-A or rTGF-β1 in the indicated concentrations. Cells were then plated in glucose-free media containing rIL-15 at 5 ng/mL for 3 h, followed by transfer to gelatin-coated seahorse assay plates for measurement of extracellular acidification rate (ECAR, C) and oxygen consumption rate (OCR, D) every 7 min using the Seahorse XF^e^96 analyzer. Data represented as mean ± SEM of three biological replicates and paired T test was used for statistical analysis, where ***p < 0.001 for comparisons between the indicated stimulation within WT or RII^FL^ groups, and ^###^p < 0.001 for comparisons between WT and RII^FL^ cells.

To relate these changes to cellular metabolism, we also measured the extracellular acidification rate (ECAR) and oxygen consumption rate (OCR) of WT and TgfbRII-deficient NK cells in response to rActivin-A or TGF-β (fig 4C-D). Surprisingly, TgfbRII-deficient NK cells displayed at least 2-fold higher basal glycolysis (ECAR) and oxidative phosphorylation (OCR) compared to WT cells suggesting they have an enhanced metabolism *in vitro*. Although rTGF-β1 stimulation was unable to inhibit ECAR and OCR in TgfbRII-deficient NK cells, it completely suppressed WT NK cell metabolism. rActivin-A inhibited both these parameters in both WT as well as in TgfbRII-deficient NK cells and this effect was saturated at the lower concentration of rActivin-A suggesting minimal SMAD2/3 phosphorylation is required to suppress metabolism (fig. 4C-D). Taken together, these results demonstrate for the first time that the ALK4 pathway regulates the cellular metabolism in murine NK cells.

### Activin-A modulates NK cell killing

rTGF-β1 has been shown to abolish NK cell cytotoxicity *in vitro* (Viel et al., 2016), and likewise the constitutive activation of TGF-βR1 in NKp46^+^ cells completely abolishes the *in vivo* killing potential of NK cells (Gao et al., 2017). In addition to being a critical survival factor (Huntington et al., 2007; Sathe et al., 2014), IL-15 induces the expression of Granzyme B (GrzB) in NK cells (Fehniger et al., 2007). To test whether Activin signaling also limits NK cell cytotoxicity, WT or TgfbRII-deficient NK cells were cultured in IL-15 and exposed to either TGF-β or Activin. As expected, rTGF-β1 efficiently downregulated GrzB expression in WT NK cells, while rActivin-A modestly downregulated its expression in both WT and TgfbRII-deficient NK cells (fig. 5A-B). In line with this observation, we found that rTGF-β1 could suppress NK-mediated killing of B16F10 melanoma cells to a greater extent than rActivin-A (fig. 5C). These results show a subtle, but novel role for Activin in suppressing NK cell cytotoxicity, independently of canonical TGF-β signaling.

**Figure 5.**
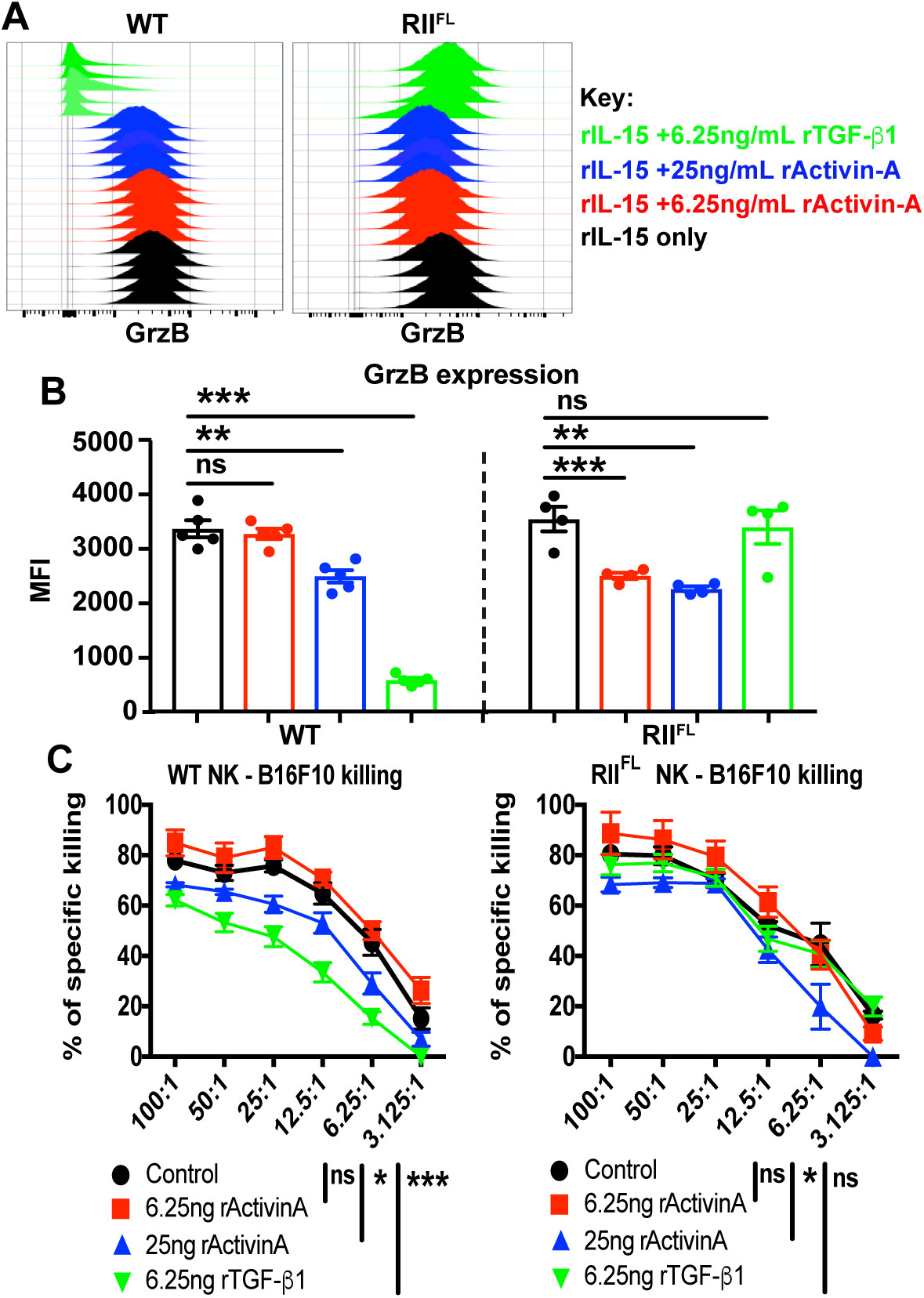
Effects of Activin signaling on NK cell cytotoxicity. (A-B) NK cells sorted from spleens of WT, or RII^FL^ NK cells were cultured *in vitro* for 5 days in 50 ng/mL rIL-15 with or without indicated concentrations of rActivin-A, or TGF-β1. At endpoint, cells were stained with intracellularly with anti-GrzB and analyzed by flow cytometry. (A) Each overlay represents one independent biological replicate of the indicated cell genotype for intracellular GrzB staining in the indicated treatment groups. (B) Quantification of GrzB stain by mean of florescence intensity (MFI) comparison of data displayed in (A). Results are expressed as the mean + SEM, each data point represents one independent biological replicate and one-way ANOVA was used for comparative statistical analysis, where *p < 0.05, **p < 0.01. (C) Sorted NK cells were cultured overnight (16 h) in presence of 50 ng/mL rIL-15 with or without indicated concentrations of rActivin-A, or TGF-β1 in animal free media, and then tested against B16F10 target cells at various effector:target ratios in a 4 h calcein dye release assay. Results are expressed as the mean ± SEM, and two-way ANOVA was used for comparative statistical analysis where *p < 0.05, and ***p < 0.001

### Distinct effects of Activin and TGF-β signaling pathways on cytokine secretion by NK cells

To further assess if Activin and TGF-β signaling pathways have divergent impacts on NK cell function, cytokines were screened from the supernatant of purified WT or TgfbRII^fl/fl^ NK cells after stimulation with rIL-15 with or without rActivin-A, rTGF-β1, rIL-12 and rIL-18 for 48 h. Measurement of secreted cytokines revealed that TGF-β signaling impairs GM-CSF production in response to IL-12 and IL-18 stimulation, whereas Activin-A signaling failed to impact on GM-CSF production (fig. 6A). IFN-γ production by NK cells is dramatically increased in response to IL-12 and IL-18 stimulation and this was not affected by either Activin-A or TGF-β in this setting. Interestingly, despite being phenotypically similar to WT NK cells, TGF-βR-deficient NK cells produced significantly more IFN-γ than WT cells suggesting they are primed in the absence of TGF-β signaling *in vivo* (fig. 6B). To test whether these findings could be corroborated with human NK cells, cord blood-derived NK cells were cultured for 48 h in the same concentrations of IL12, IL-15 and IL-18 as mouse cells, and in a range of Activin-A or TGF-β concentrations. All TGF-β1 concentrations (6.25, 25, and 100 ng/mL), were able to suppress human IFNg secretion following IL-12 and IL-18 stimulation, whilst only the highest concentration of Activin-A (100 ng/mL), achieved this effect (fig. 6C). This result supports a previous report (Robson et al., 2009) which demonstrated similar effects in human peripheral blood-derived NK cells, but also reveals that not all aspects of Activin-A or TGF-β signaling are conserved between mouse and human cells.

**Figure 6.**
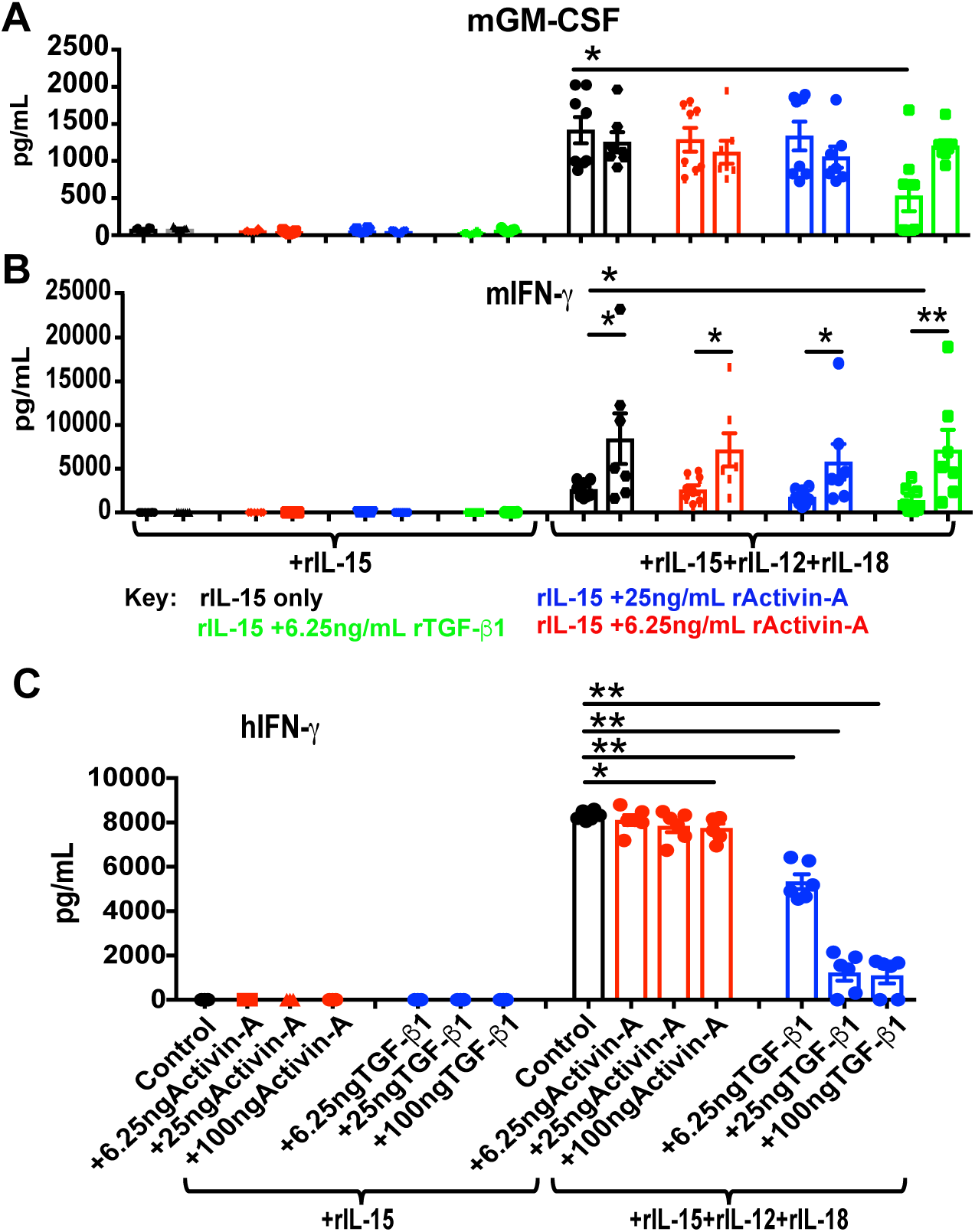
Differential cytokine induction or suppression by Activin-A and TGF-β signaling in NK cells. (A-B) NK cells sorted from spleens of WT (data displayed in the first column of each group), or RII^FL^ (data displayed in the second column of each group) mice were cultured in 50 ng/mL rIL-15, with or without 20 pg/mL rIL-12 and 50 ng/mL rIL-18, with or without the indicated concentrations of rActivin-A or rTGF-β1 for 48 h. Supernatants were then analyzed for secretion of murine GM-CSF (A) or IFN-γ (B). (C) Human cord blood NK cells were isolated by negative selection and cultured in 50 ng/mL rIL-15, with or without 20 pg/mL rIL-12 and 50 ng/mL rIL-18, with or without the indicated concentrations of rActivin-A or rTGF-β1 for 48 h. Results are expressed pg/mL (y axis) as the mean + SEM. Each data point represents one independent biological replicate and unpaired T test was used for comparative statistical analysis, where *p < 0.05.

### Targeted inhibition of Activin signaling in NK cell-dependent in vivo models

Using clinical grade Follistatin (98% homologous between mouse and human), we first tested whether it was possible to therapeutically enhance the highly NK cell-dependent immune response against the B16F10 melanoma model (Sathe et al., 2014; Souza-Fonseca-Guimaraes et al., 2015). Follistatin was administered according to the manufacturer’s suggestions (Paranta Biosciences Ltd, Melbourne, VIC, Australia), and compared to a TGF-β1/2/3-neutralizing antibody (Clone 1D11) previously validated for *in vivo* use (Gartlan et al., 2015). The data revealed that similar to TGF-β blockade, Follistatin significantly reduced the number of lung metastases, but no additive benefit was observed when both agents were combined (fig. 7A). In a different approach, we compared the effect of Follistatin in a higher tumor burden assay by intravenously injecting the double of B16F10 cells (4×10^5^) in RII^FL^ mice or the respectively wild type littermate control and found that only Follistatin treatment significantly reduced lung metastases with more enhanced effect when TgfbRII was conditionally deleted in NK cells (RII^FL^ mice) (fig. 7B). These results suggested that *in vivo* inhibition of Activin-A could improve tumor rejection by enhancing the innate effector functions of NK cells.

**Figure 7.**
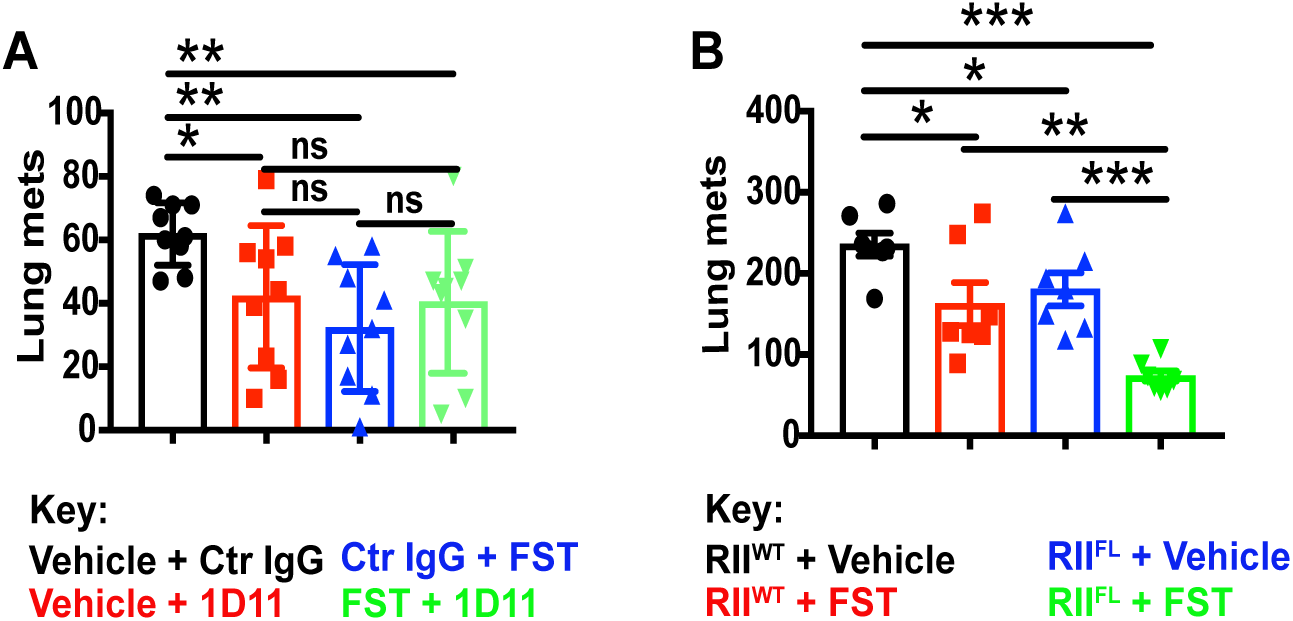
Inhibition of Activin-A or TGF-β signaling *in vivo* results in therapeutic benefit. (A) Groups of 9 WT mice were injected i.v. with 2×10^5^ B16F10 melanoma cells. Treatment regimen with Follistatin, or FST (10 µg i.p. per mouse per day, starting 1-day post tumor inoculation), and neutralizing anti-TGF-β1,2,3 antibody (500 µg i.p. dose per mouse every two days, starting 1 day post tumor inoculation) was administered until endpoint (14 days post tumor inoculation). (B) Groups of 6-7 (RII^FL^) or respective littermate controls mice (RII^WT^) were injected i.v. with 4×10^5^ B16F10 cells. Treatment regimen with FST and experimental endpoint was performed as described for (A). Symbols in scatter plots represent the number of tumor colonies in the lung from individual mice (with mean and SEM shown by cross-bar and errors. Mann-Whitney test was used to compare differences between groups of mice as indicated, where *P <0.05, **P < 0.01 and ***P < 0.01 were considered for statistical significance.

TGF-β signaling in NK cells was recently shown (Cortez et al., 2016; Gao et al., 2017) to induce cellular transdifferentiation from NK cells into ILC1-like cells *in vivo*. Although the conditional genetic deletion of TGF-βRII in NKp46^+^ cells suggested that this transdifferentiation could be largely prevented *in vivo*, alternate stimuli of the SMAD2,3 pathway likely compensated for the lack of TGF-β signaling. Murine cytomegalovirus (MCMV) infection was previously described to induce the upregulation of ILC1-like molecules (e.g. TRAIL, CD49a, CD69) (Schuster et al., 2014) on salivary gland-NK cells in a TGF-β-like manner (Cortez et al., 2016). To assess if this MCMV-related transdifferentiation could be therapeutically prevented, we treated MCMV-infected mice with Follistatin, a TGF-β-neutralizing antibody, or a combination of both. Notably, we did not observe any significant differences in viral titers in the lungs, salivary glands, or spleens the different treatment groups at day 6 days post-infection (fig. S6). Similarly, neither treatment prevented the upregulation of ILC1-like markers (e.g. CD49a, CD69, and TRAIL) on NKp46^+^ cells in the liver or salivary glands on day 6 post infection (fig. S7). These data show that the MCMV replication and expression of NK cell surface markers are not affected by the therapeutic inhibition of Activin-A or TGF-β1 during acute infection.

### Validation of Activin signaling in human NK cells

To further validate whether Activin-A signaling is conserved between human and mouse we examined key signal transduction pathways and markers of NK cell transdifferentiation. Human cord blood-derived NK cells were purified and activated in media containing donor-matched serum, IL-15 and a m/hTGF-β1/2/3-neutralizing antibody was used for 5 days to remove any TGF-β present in the serum. Cells were then exposed to either rIL-15, rActivin-A, rTGF-β1 or combinations of these cytokines for the time periods indicated. Assessment of the protein phosphorylation of signaling molecules revealed efficient phosphorylation of SMAD2/3 by either 100 ng/mL rActivin-A or rTGF-β1, irrespective of whether IL-15 was present (fig. 8A and fig. S8A). In agreement with our previous report (Viel et al., 2016), we observed that IL-15-induced phosphorylation of STAT5 is independent of SMAD2,3 phosphorylation (either by Activin-A or TGF-β1) (fig. 8B and fig. S8B). However, Activin-A or TGF-β1 were able to suppress the IL-15-induced phosphorylation of AKT and S6K controlled metabolic pathways (fig. 8C-D, and fig. S8C-D, respectively), in line with data in (fig. 4C-D) and our previous work on mouse NK cell metabolism (Viel et al., 2016).

**Figure 8.**
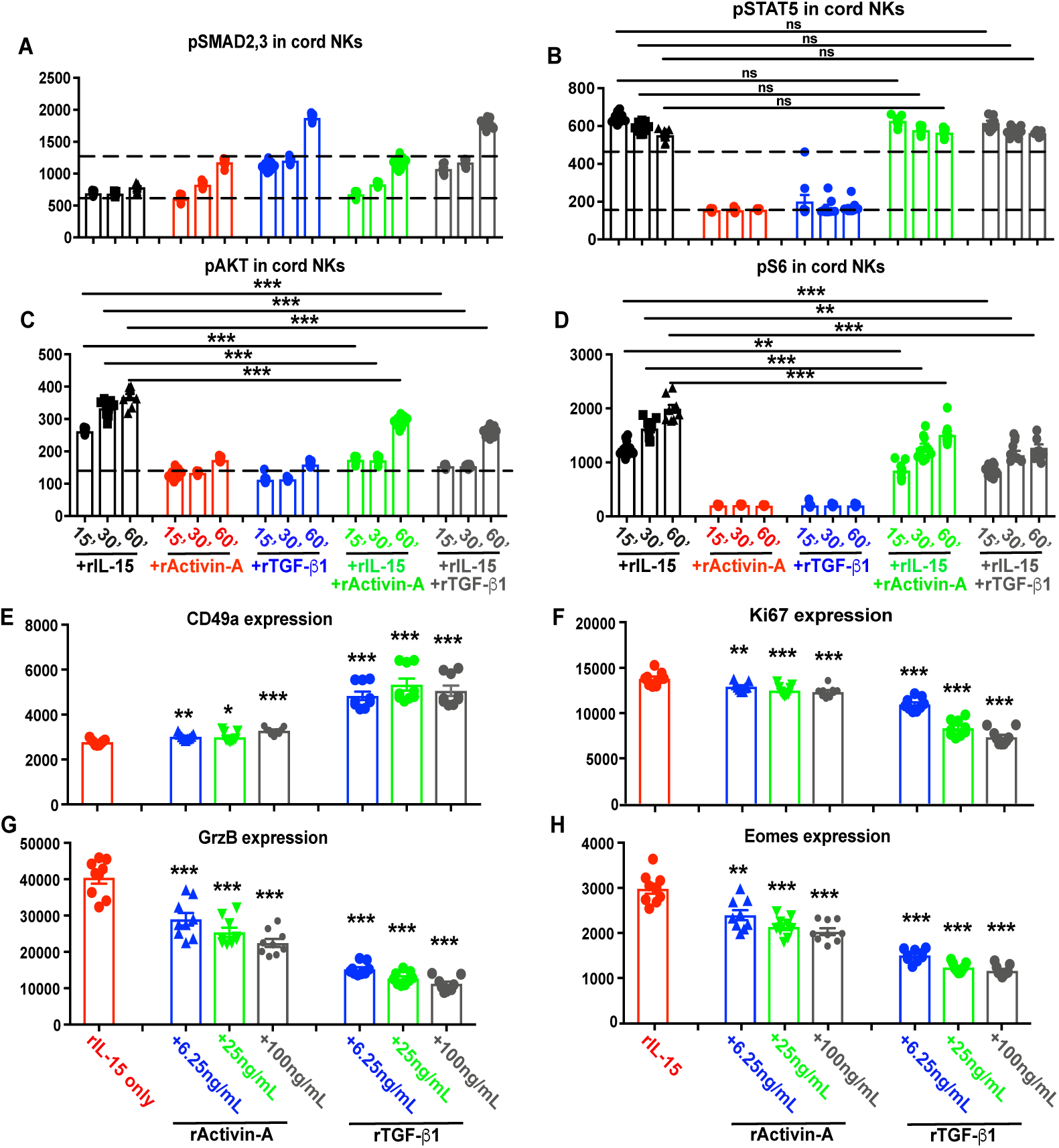
Conserved features of Activin signaling in human NK cells. (A-H) NK cells were from fresh cord blood of 3 different donors and primed in rIL-15 for 48 h. Cells were then washed, starved for 4 h in animal component-free media at 37°C, and stimulated in rIL-15 media (10 ng/mL) with indicated concentrations of rActivin-A or rTGF-β1 for 1 h. p SMAD2,3 (A), pSTAT5 (B), pAKT (C) and pS6 (D) were compared by flow cytometry as MFI. E-H: A portion of the same cord blood NK samples, were separately cultured for 7 days in presence of 50 ng/mL of rIL-15 with or without the indicated concentrations of rActivin-A or rTGF-β1. At endpoint, cells were cell surface stained with anti-CD49a (E), intracellularly stained with and anti-Ki67 (F), anti-GrzB (G) and anti-Eomes (H) antibodies and its MFIs were analyzed by flow cytometry. Results are expressed as the mean ± SEM, each graph dot represents one independent technical replicate (with 3 technical replicates for each donor (3)) and unpaired T test was used for comparative statistical analysis, where *p < 0.05, **p < 0.01 and ***p < 0.001.

Finally, we also examined whether Activin-A signaling could induce the upregulation of ILC1 markers on human NK cells similar to what we observed in mouse NK cells (fig. 3). For this, human cord blood-derived NK cells were cultured for 7 days in IL-15 with either rActivin-A and rTGF-β1. Consistent with our findings using murine cells, we detected upregulation of CD49a in response to both rActivin-A and rTGF-β1 in a concentration-dependent manner (fig. 8E and fig. S8E) and a downregulation of GrzB and Ki67 expression (fig. 8F-G and fig. S8F-G). Differences were also observed with the murine findings, as rActivin-A was enough to downregulate Eomes expression in human NK cells (fig. 8H and fig. S8H). Finally, similar to our findings in murine NK cells (fig 1F-G), however rActivin-A or rTGF-β1 both reduced the expansion of human NK cells by suppressing their proliferation rate (reduced mean division number) (fig. 9A-B) without affecting their survival (data not shown). These findings reveal that the regulation of NK cell biology by the ALK4 signaling pathway is conserved between humans and mice.

**Fig. 9.**
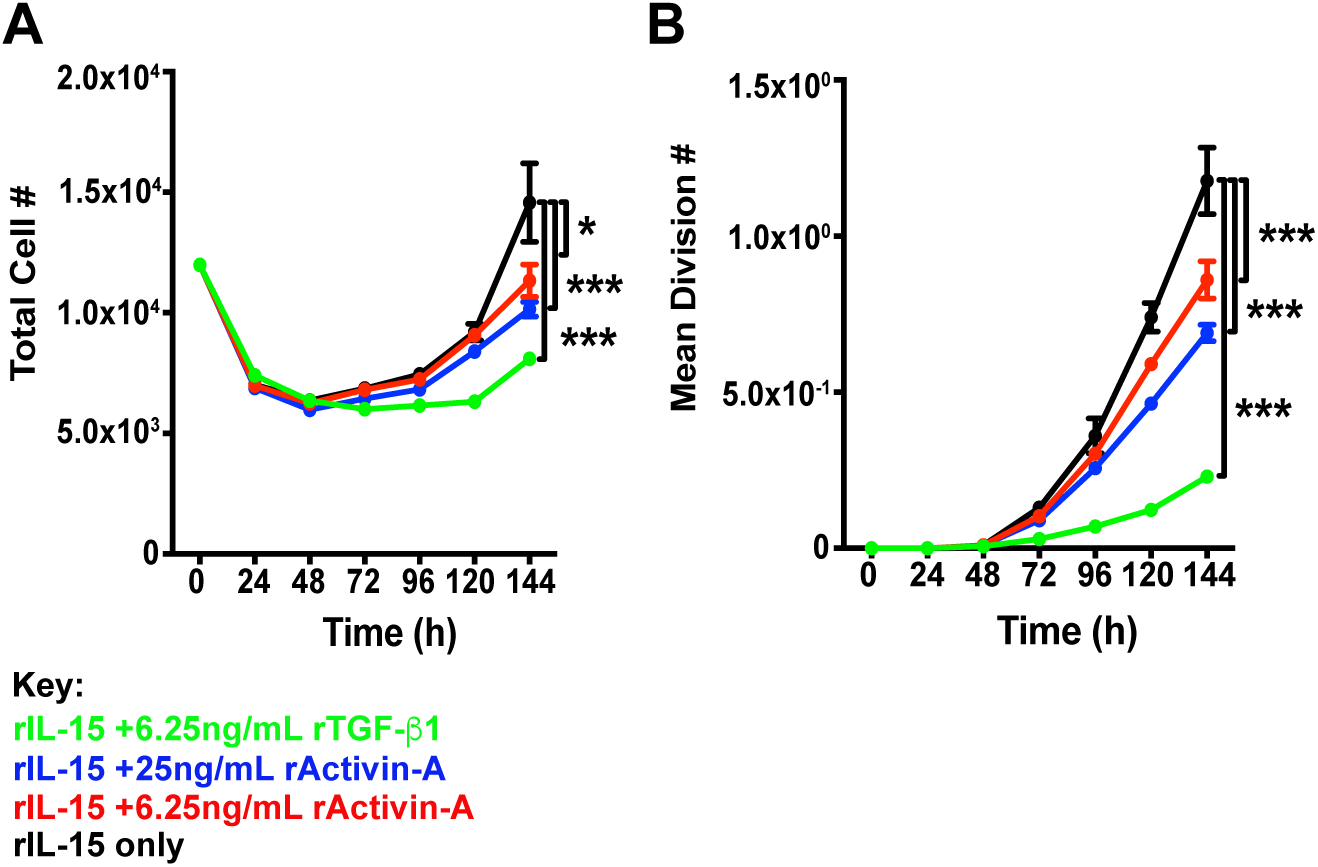
Activin signaling impairs human NK cell proliferation. (A-B) Freshly-isolated human cord blood-derived NK cells were labelled with a division-tracking dye and cultured in the presence of 50 ng/mL rIL-15 in addition to rTGF-β1 or rActivin-A as indicated. For each condition, the total number of live cells was enumerated at 24 h intervals (A), and the population average division number was calculated (B). Results shown are representative from one donor, of 3 independent biological replicates. Error bars represent the S.D. of 3 technical replicates, and two-way ANOVA tests were used for comparative statistical analysis where *p < 0.05 and ***p < 0.001.

## Discussion

Immune “checkpoint” inhibitors have revolutionized cancer therapy by reactivating tumor-resident cytotoxic lymphocytes to kill the malignant cells comprising the tumor mass. NK cells possess an innate ability to detect cellular transformation and are key to cancer immunosurveillance, particularly in the prevention of metastasis (Krasnova et al., 2017). An understanding of the tumor microenvironment and how tumor cells evade detection by NK cell is now emerging and has stimulating great interest in therapeutic targeting of such pathways. As detailed below, we recently revealed a novel immune evasion mechanism in which tumors exploit the TGF-β signaling pathway to transdifferentiate conventional NK cells (CD49a^neg^) into ILC1-like subsets (CD49a^+^), and in doing so, lower their intrinsic anti-tumor functions (Gao et al., 2017). An earlier study described a similar transdifferentiation in salivary gland NK cells (Cortez et al., 2016). Both of these studies used the conditional deletion of TGF-βRII gene in NK cells (NKp46^cre/+^TgfbRII^fl/fl^) as negative control for TGF-β signaling in NK cells. Interestingly, despite NKp46^cre/+^TgfbRII^fl/fl^ NK cells being entirely unresponsive to recombinant TGF-β1 *in vitro*, both studies displayed a minor ILC1-like NKp46^+^CD49a^+^ population *in vivo* in NKp46^cre/+^TgfbRII^fl/fl^ mice. Together, this suggests that factors other than TGF-β1 might also mediate NK cell reprograming and tumor immune evasion.

Our recent work on TGF-β-dependent immune evasion mechanism by effector cNK cells (Eomes^+^, CD49b^+^, CD49a^neg^, TRAIL^neg^) transdifferentiation into ILC1-like cells (Eomes^neg^, CD49b^neg^, CD49a^+^, TRAIL^+^) or intermediate (int)ILC1 (Eomes^+^, CD49b^+^, CD49a^+^, TRAIL^+^) has been observed in experimental metastasis and orthotopic melanoma models, as well as for fibrosarcomas, suggesting that TGF-β can suppress both systemic and tumor-resident NK cell response (Gao et al., 2017). Given our data demonstrates that Activin-A acts in a similar manner to TGF-β in inducing an ILC1-like phenotype on NK cells, this may help explain the residual ILC1-like cells observed in tumors of NKp46^cre/+^TgfbRII^fl/fl^ mice. Our *in vitro* data clearly demonstrate that Activin-A induces several changes to murine or human NK cells reminiscent of TGF-β. These include suppression of cellular metabolism and proliferation, upregulation of CD69, CD49a and TRAIL, and the downregulation of Eomes, Ki67 and Granzyme B and Perforin. Importantly, these changes occurred with TGF-βRII-deficient NK cells ruling out any role for Activin-A synergizing with TGF-β found in serum-containing culture media. The effect of TGF-β concentrations (~60 pg/mL) found in culture media is noteworthy as TGF-βRII-deficient NK cells consistently appear hyper-active compared to WT NK cells expanded in IL-15 *in vitro*. For example, TGF-βRII-deficient NK cells displayed significantly higher basal and maximal metabolism and produce significantly more IFN-γ compared to WT counterparts. Interestingly, while Activin-A was able to suppress TGF-βRII-deficient NK cell metabolism, it only impaired glycolysis and oxidative phosphorylation to levels equivalent with WT NK cells cultured in the absence of Activin-A highlighting the enhanced metabolic state of TGF-βRII-deficient NK cells.

The concentration-dependent effect of Activin-A on WT NK cells was exemplified in experiments examining SMAD2/3 phosphorylation, Ki67 expression, Granzyme B expression and impaired B16F10 killing levels. However, NK cell metabolism and ILC1 gene induction appeared to be maximally altered at the lowest concentrations of Activin-A, highlighting the differential sensitivity of certain NK-suppressive pathways to Activin-A signaling. Despite this, endogenous levels of Activin-A appeared sufficient to suppress NK cell activity as the therapeutic administration of Follistatin reduced lung metastases in a NK cell-dependent melanoma model. Taken together, our results reveal a novel capacity for Activin-A to regulate NK transdifferentiation, thereby ultimately facilitating tumor evasion of NK cell immune surveillance in a TGF-β independent mechanism. This study suggests that combinatorial therapies targeting Activin signaling may result in greater prevention of cNK transdifferentiation in the tumor microenvironment (and associated suppression of cellular metabolism and effector function) and subsequently enhance innate anti-tumor immune control.

## Materials and Methods

### Mice

TgfbRII-deficient NK cells were isolated from a NKp46^cre/wt^TgfbRII^fl/fl^ mice, obtained by crossing the NKp46-iCre (Narni-Mancinelli et al., 2011) with the TgfbRII LoxP mice (Leveen et al., 2002) as previous described by our group (Gao et al., 2017; Viel et al., 2016). WT NK cells were isolated from the respectively littermate controls (NKp46^wt/wt^TgfbRII^fl/fl^). All mice were on a C57BL/6J background, bred and maintained at the Walter and Eliza Hall Institute of Medical Research (WEHI). For murine cytomegalovirus infection, BALB/C mice were acquired from the Animal Research Centre (ARC, Perth) and housed in the Bioresources Department of the Harry Perkins. All experiments were performed using cells from age/sex matched cohort of mice (age range 8-12 weeks). Cohort sizes are described in each figure legend to achieve statistical significance. No biological replicate was excluded based on pre-established criteria. All experiments were approved by the WEHI and Harry Perkins Institute’s Animal Ethics Committees.

## Reagents

Reagents or antibodies targeting the following human (h) or murine (m) epitopes were purchased from Biolegend (San Diego, CA): 7-AAD, mCD3 (145-2C11), mCD19 (6D5), mCD49b (DX5 and HMα2), mCD62L (MEL-14), mDNAM-1 (10E5), mF4/80 (BM8), mLy6G (1A8), mNK1.1 (PK136), mNKp46 (29A1.4), Streptavidin-FITC, and mTCR-β (H57-597). Reagents or antibodies targeting the following epitopes were purchased from eBioscience (San Diego, CA): mCD49b (DX5), hEomes (WD1928), mEomes (Dan11mag), mGranzyme B (NGZB), hGranzyme B(GB11), mTCR-β (H57-597), and mTRAIL (N2B2). Reagents or antibodies targeting the following epitopes were purchased from BD Biosciences (San Jose, CA): Akt(pS476) (M89-61), Fixable Viability Stain, mNK1.1 (PK136), Ki67 (B56), SMAD2(pS465/S467)/SMAD3(pS423/pS425) (I72-670) and STAT5(pY694) (47/Stat5(pY694)). Antibody targeting pS6(pS235/S236) (#2211) was purchased from Cell Signaling Technology (Danvers, MA, USA). Antibodies targeting mCD45.2 (30F11), mCD49a (REA493), hCD49a (TS2/7), mCD69 (H1.2F3) and hNKp46 (9E2) were purchased from Miltenyi Biotec (Bergisch Gladbach, Germany). MACSXpress Human NK cell Isolation Kit, was used for negative isolation of cord blood NK cells (Miltenyi Biotec). For intracellular Eomes, granzyme B and Ki67 detection, cell surface stained samples were fixed and permeabilized using Intracellular Fixation & Permeabilization Buffer Set and stained with antibodies in 1X Permeabilization Buffer (eBioscience). Cell number was calculated by using BD Liquid Counting Beads (BD Biosciences).

### NK cell isolation and culture conditions

Mice were sacrificed and spleens were harvested and prepared for flow cytometry as previously described. Spleen homogenates were incubated in Fc blocking buffer (2.4G2 antibody) on ice for 15 min and then pre-enriched by lineage (Lin: CD3, CD19, CD49a, F4-80, Ly6G, TCRβ)-biotin antibody cocktail staining followed by MagniSort Streptavidin Bead Negative Selection (eBioscience). The remained fraction was stained with Streptavidin-FITC and NK cells (7-AAD^-^Lin^-^CD45^+^NK1.1^+^NKp46^+^CD49b^+^), from indicated transgenic or WT mice, were then sorted by BD FACS Aria II cell sorter (BD Biosciences) to achieve a final cell purity of 99-100%. Human cord blood NK cells were isolated by negative selection with a final cell purity of 95-99%, using the MACSXpress Human NK Cell Isolation Kit (Miltenyi Biotec). Following sorting/negative selection, mouse or human NK cells were stained with Cell Trace Violet (InVitroGen), and plated at a density of 25,000 cells per well in V bottom 96-well plates (Greiner Bio-One, Frickenhausen, Germany) containing RPMI 1640 supplemented with 10% FCS, 1% sodium pyruvate (Gibco, Grand Island, NY), 1% Glutamax (Gibco), 10 mM HEPES, 0.1% 2-mercaptoethanol (Gibco), 1% penicillin/streptomycin in presence of the indicated concentrations, and time points, of recombinant human rIL-15 (Miltenyi Biotec), recombinant human/mouse Activin-A (R&D Systems, Minneapolis, MN), recombinant mouse TGF-β1 (eBiosciences) or recombinant human TGF-β1 (Peprotec, Rocky Hill, NJ). Culture conditions were maintained at 37°C and 5% CO_2_. Intracellular staining of phosphorylated signaling proteins was performed using anti-pAkt, pSMAD2,3 antibody, anti-pS6 and anti-pSTAT5 after endpoint fixation and permeabilization with Lyse/Fix and Perm III buffers (BD Biosciences). Data was acquired using LSR Fortessa Flow Cytometer (BD Biosciences). Flow cytometric analysis was performed using Flowjo (Treestar, Ashland, OR) software. For proliferation and viability assays, fresh human and murine NK cells were incubated with 5 µM cell trace violet (Thermo Fisher Scientific) according to the manufacturer’s instructions, and 8 x 10^3^ labeled cells were seeded into 96-well round bottom plates in complete media with the indicated cytokine concentrations and time points. Routine time points were assessed on a BD FACS Verse cytometer (BD Biosciences), and survival and division numbers were determined using the precursor cohort-based method (Hawkins et al., 2007; Marchingo et al., 2014), as previously described for NK cell kinetics by our group (Viant et al., 2017).

### In vitro cytokine secretion assays

FACS Aria-sorted NK cells from the spleen (Lin^neg,^ 7AAD^neg^, CD49a^neg^, NKp46^+^, NK1.1^+^, CD49b^+^) of the indicated mouse genotypes, were expanded for up to 10 days in complete RPMI 1640 containing 10% FBS, 1 µg/ml of anti-TGF-β1,2,3 blocking antibody (clone 1D11.16.8 from Bio X Cell, West Lebanon, NH), β-Me, GlutaMax and sodium pyruvate (Gibco). For cytokine secretion assays, cells were stimulated in the indicated concentrations of rTGF-β1, rActivin-A, or rIL-15 culture conditions in animal free / TGF-β1 free TexMACS medium (Miltenyi Biotec) containing β-Me, GlutaMax and sodium pyruvate for 48 hours. In certain proliferation assays, the pan adenosine receptor agonist 5’-(N-Ethylcarboxamido) adenosine (NECA) (Sigma Aldrich, St Louis. MO) was used at 1 µM in the culture media for comparison with TGF-β1. For the detection of cytokines in the supernatants of *in vitro* assays, IFN-γ was measured by ELISA with the respective human or murine IFN-γ Duoset Kit (R&D Systems) according to the manufacturer’s instructions, while all other indicated cytokines were detected using Cytometric Bead Array (CBA) technology (BD Biosciences) according to the manufacturer’s instructions.

### Target : effector cell co-cultures

Sorted and expanded NK cells (as described above) were used to perform standard 4 hours cytotoxicity assays using Calcein-AM (BD Biosciences)-labelled target cells (B16F10 melanoma cells) as previously described. Briefly, NK cells from the indicated genotypes were cultured overnight (16 h) in the presence of rTGF-β1 or rActivin-A in 50 ng/mL of rIL-15 at the indicated concentrations. NK cells were then seeded at the indicated ratio with Calcein-AM-labelled target cells in complete NK cell medium (phenol-red free RPMI 1640 containing 10% FBS, β-Me, GlutaMax and sodium pyruvate, all from Gibco). After 4 hours co-culture, supernatants were transferred to opaque 96 well-plates (Costar, Corning, NY) and fluorescence emission was measured (excitation filter: 485 ± 9nm; cut off: 515 nm; emission 525 ± 15nm) using a SpectraMax M5e microplate reader (Molecular Devices, Sunnyvale, CA). Cytotoxicity data was expressed as percent lysis relative to the spontaneous (target cells alone) and maximum release (1% Triton X-100, Sigma Aldrich, treated cells). In some experiments, melanoma cells were cultured in round glass coverslips and co-cultured overnight in 24-well plates with purified NK cells in the indicated culture conditions. Cells derived from the different assayed conditions were processed for scanning electron microscopy (SEM). In summary, coverslips-containing cells were fixed in a 24-well plate with 2.5% glutaraldehyde (0.1 M cacodylate buffer, pH 7.2), washed with PBS and post-fixed in 1% OsO_4_ for 30 min in the dark at room temperature. After washing, the cells were dehydrated using increasing ultra-pure ethanol concentrations. All reagents for electron microscopy were purchased from Electron Microscopy Sciences (Fort Washington, PA). Coverslips were dehydrated using CO_2_ critical point, metalized and cells were analyzed using a JEOL JSM-6360 LV scanning electron microscope (JEOL, Peabody, MA).

### NK cell preparation for proteomics

NK cells were sorted from the spleen of RII^FL^ (*n* = 3) or control mice (*n* = 4), expanded for 7 days in 50 ng/mL of rIL-15 in RPMI 1640 media + 10% FCS supplemented with neutralizing TGF-β1,2,3 antibody (1D11.16.8). Cells were then washed 3 times with cold PBS, starved in serum free media for 4 hours, and then 5×10^6^ cells per replicate/culture condition were cultured with rIL-15 (10 ng/mL) with rTGF-β1 (6.25 ng/mL) for 24 hours. Cells were then washed three times with ice cold PBS prior to dry cell pellet storage at -80 **°**C. Cells were lysed in KALB lysis buffer supplemented with protease inhibitors (Complete Cocktail tablets, Roche), 1 mM PMSF, 1 mM Na_3_VO_4_ and 1 mM NaF and incubated for 1 h on ice. Lysates were clarified by centrifugation at 14,000 g for 15 min at 4 **°**C. Protein concentrations were determined using the BCA method (Pierce, Rockford). NK cell protein lysates (50 µg) were prepared for mass spectrometry analysis as described by Hughes *et al.* (Hughes et al., 2014) with modifications. For all our experiments with magnetic beads, we used a 1:1 combination mix of the two types of commercially available carboxylate beads (Sera-Mag Speed beads, #24152105050250, #44152105050250, Thermo Fisher Scientific). Beads were prepared freshly each time by rinsing with water three times prior to use and stored at 4**°**C at a stock concentration of 10 µg/µL. Cell lysates were reduced with 2 M Dithiothreitol (DTT, 50 mM final conc.) for 1 h at 37 **°**C.

Samples were then alkylated with 1M Iodoacetamide (IAM) (100 mM final conc.) for 30 mins in the dark at room temperature (RT). Samples were quenched with 2M DTT (250 mM final conc.). Lysates were acidified with formic acid (FA, pH <3) and 7 µl of the concentrated bead stock carboxylate beads (10 µg/µL) was added. Acetonitrile (ACN) was added to reach a final concentration of 50% (v/v). Mixtures were left to incubate upright at RT for 1 h. Pre-chilled (−20 **°**C) acetone/0.1% FA was added to the protein/bead mixtures at a 4x volume (v/v) and left for 20 min at -20 **°**C to allow proteins to precipitate onto the beads. The beads were washed twice with pre-chilled 80% acetone/1% FA (250 µL washes). A further 100 µL 80% acetone/1% FA was used to transfer the bead/acetone suspension into a PCR plate for subsequent enzymatic digestion. Acetone was completely evaporated from the PCR plate prior to the addition of 40 µL digestion buffer (10% 2-2-2-Trifluorethanol (TFE)/100 mM NH_4_HCO_3_) containing 2 µg Trypsin-gold (Promega, V5280) at a 1:25 enzyme: substrate ratio. The plate was briefly sonicated in a water bath to disperse the beads, and the plate transferred to a ThermoMixer instrument for digestion at 37 **°**C for 1 h (1200 rpm). The supernatant comprising of peptides was then collected from the beads using the magnetic rack and an additional elution (20 µL 2% Dimethyl sulfoxide, DMSO, Sigma) was performed on the beads. The eluates were pooled together and transferred to pre-prepared C8 stage tips to ensure no beads are carried through the analysis. Two plugs of C8 resin (Empore, 2214) were prepared in 200 uL un-filtered tips, pre-wetted with 50 µL ACN. The pooled peptides were then added to the spin tip and the eluate collected into a fresh lo-bind Eppendorf tube. An additional 50 µL of 50% ACN/0.1% FA was added to the spin tip and collected into the same tube. The eluates were lyophilized to dryness using a CentriVap (Labconco) prior to reconstituting in 25 µL Buffer A (0.1% FA/2% ACN) ready for MS analysis.

### Mass spectrometry analysis

Peptides (2 µL) were separated by reverse-phase chromatography on a 1.9 µm C18 fused silica column (I.D. 75 µm, O.D. 360 µm x 25 cm length) packed into an emitter tip (Ion Opticks, Australia), using a nano-flow HPLC (M-class, Waters). The HPLC was coupled to an Impact II UHR-QqTOF mass spectrometer (Bruker, Bremen, Germany) using a CaptiveSpray source and nanoBooster at 0.20 Bar using acetonitrile. Peptides were loaded directly onto the column at a constant flow rate of 400 nL/min with buffer A (99.9% Milli-Q water, 0.1% formic acid) and eluted with a 90 min linear gradient from 2 to 34% buffer B (99.9% acetonitrile, 0.1% formic acid). Mass spectra were acquired in a data-dependent manner including an automatic switch between MS and MS/MS scans using a 1.5 second duty cycle and 4 Hz MS1 spectra rate followed by MS/MS scans at 8-20 Hz dependent on precursor intensity for the remainder of the cycle. MS spectra were acquired between a mass range of 200–2000 m/z. Peptide fragmentation was performed using collision-induced dissociation (CID).

For data analysis, raw files consisting of high-resolution MS/MS spectra were processed with MaxQuant (version 1.5.8.3) for feature detection and protein identification using the Andromeda search engine (Cox et al., 2011). Extracted peak lists were searched against the *Mus musculus* database (UniProt, October 2016), as well as a separate reverse decoy database to empirically assess the false discovery rate (FDR) using strict trypsin specificity allowing up to 2 missed cleavages. The minimum required peptide length was set to 7 amino acids. In the main search, precursor mass tolerance was 0.006 Da and fragment mass tolerance was 40 ppm. The search included variable modifications of oxidation (methionine), amino-terminal acetylation, the addition of pyroglutamate (at N-termini of glutamate and glutamine) and a fixed modification of carbamidomethyl (cysteine). The “match between runs” option in MaxQuant was used to transfer identifications made between runs within a group of samples on the basis of matching precursors with high mass accuracy (Cox and Mann, 2008). PSM and protein identifications were filtered using a target-decoy approach at an FDR of 1%.

### RNA sequencing and bioinformatics analysis

From the same cells in the respectively culture conditions detailed in “NK cell preparation for proteomics”, 10^6^ cells per replicate or culture condition were prepared for RNA sequencing. Total RNA was extracted using RNAEasy Mini Kit Plus (Qiagen). Quantity of the total RNA was checked using Qubit RNA HS (Thermo Fisher Scientific). 500 ng total RNA was used for library preparation according to standard protocols (QuantSeq 3’ mRNA-Seq FWD, Lexogen). Indexed libraries were pooled and sequenced on a NextSeq500 (Illumina). 5-15 million single-end 75bp reads were generated per sample. 3’RNA-seq Single-End reads up to 75 bp were processed using *fastqc* (available online at: http://www.bioinformatics.babraham.ac.uk/projects/fastqc) and *trimmomatic* (Bolger et al., 2014) to assess read quality and trim any bases with sequencing quality less than 20 and the Illumina adapters from the end of the reads. Only reads larger than 20bp in length after trimming were kept. Reads were aligned to mm10 using *subread* (Liao et al., 2013). Duplicate reads in the resulting BAM files were marked using Picard tools (available online at: http://broadinstitute.github.io/picard). The aligned reads were sorted and indexed using *samtools* (Li et al., 2009). Reads were summarized at the gene level using *featureCounts* (Liao et al., 2014). The differential expression analysis was done in *limma* (Ritchie et al., 2015). Pathway enrichment and gene set testing was done using *limma* and EGSEA R/Bioconductor packages (Alhamdoosh et al., 2017).

### Seahorse assays

For seahorse assays NK cells were stimulated overnight (20h) in 100 ng/mL of rIL-15 in the indicated rActivin-A and r-TGF-β1 concentrations, washed 3x in PBS, incubated for 3 hours in 5 ng/mL of rIL-15 in Seahorse XF Media unbuffered glucose-free DMEM (Seahorse Bioscience, North Billerica, MA). Stimulated cells were then then transferred to 0.5% gelatin-coated seahorse plates (Seahorse Bioscience) and suspended in 160 µL of Seahorse XF Media and then stimulated with 40 µL of same media containing a final concentration (per well) of glucose at 25 mM, oligomycin at 1 µM, FCCP at 1.5M, sodium pyruvate at 1 mM, and antimycin A at 1 µM plus rotenone at 0.1 µM. Oxygen consumption rates (OCR) and extracellular acidification rates (ECAR) were measured every 7 minutes using a Seahorse XF^e^96 analyzer (Seahorse Bioscience).

### In vivo assays

*In vivo* models were performed as previously described for i.v. inoculation of 2×10^5^ or 4×10^5^ B16F10 cells (Souza-Fonseca-Guimaraes et al., 2015), or systemic MCMV infection with 5×10^3^ PFU MCMV-K181 i.p. (Schuster et al., 2014). Treatments were performed blinded and code was revealed with witnesses at end of experiments and data analysis: Clinical grade Activin-A inhibitor, Follistatin (provided by Paranta Biosciences Ltd, Melbourne, Australia) was intraperitoneally administrated with daily doses of 10 µg per mouse as indicated; neutralizing anti-TGF-β1/2/3 antibody clone 1D11.16.8, or corresponding control IgG1 (Bio X Cell), was administrated every 3 days with intraperitoneally administered doses of 500 µg per mouse as indicated. At endpoint of MCMV infections (d6 post intraperitoneal viral inoculation), organs were processed for antibody staining and flow cytometric analysis as indicated, and MCMV viral titers in organs were determined by plaque assay using M210B4 cells as previously described (Schuster et al., 2014; Souza-Fonseca-Guimaraes et al., 2015).

### Statistics

Statistical analysis was performed using GraphPad Prism Software V6. Statistical tests used were the unpaired Mann Whitney test, paired T tests for Seahorse assays, and the Two-Way ANOVA test for cytotoxicity and proliferation experiments. Error bars represent SEM or SD, as indicated in the respective Figure Legends. Levels of statistical significance are expressed as P values: *P < 0.05, **P < 0.01, ***P < 0.001. For Label-free quantitative proteomics pipeline, statistically-relevant protein expression changes between the RII^FL^ and control groups were identified using a custom in-house designed pipeline as previously described (Delconte et al., 2016a) where quantitation was performed at the peptide level. Probability values were corrected for multiple testing using Benjamini–Hochberg method (Glueck et al., 2008). Cut-off lines with the function y= -log_10_(0.05)+c/(x-x_0_) (Keilhauer et al., 2015), were introduced to identify significantly enriched proteins. c was set to 0.2 while x0 was set to 1, representing proteins with a twofold (log2 protein ratios of 1 or more) or fourfold (log2 protein ratio of 2) change in protein expression, respectively.

## Supplementary Materials

Fig. S1. Alternative Activin receptors (ACVR2A, ACVR2B and ACVR1C) RNA expression.

Fig. S2. TGF-β1 levels in TexMACS animal free media, conventional RPMI 1646 and RPMI 1640 +10% fetal calf serum (FCS).

Fig. S3. Total NK cell cohort number after Activin-A or TGF-b treatment.

Fig. S4: Comparison of adenosine and TGF-β treatments on NK cell proliferation.

Fig. S5: Representative scanning electron micrographs of melanoma cells adhered on glass coverslips and co-cultured with sorted NK cells.

Fig. S6: MCMV viral titters after anti-TGF-β1,2,3 or Follistatin treatment.

Fig. S7: CD49a, CD69 and TRAIL expression at steady state or 6 days post MCMV infection in NKp46^+^ cells of liver or salivary gland homogenates.

Fig. S8: Conserved features of Activin signaling in human NK cells

Table. S1. Summary of the Log_2_ FC and P values associated with the label-free quantitative proteomics experiments.

## Acknowledgments

We thank all the members of the Huntington Laboratory for discussion, comments and advice on this project; Alison Campbell, Elliot Surgenor, Eren Loza, Louise Spencer, Tania Camilleri and Tobias Kratina for mouse breeding, maintenance, genotyping and technical support. **Funding:** This work is supported by project grants from the National Health and Medical Research Council (NHMRC) of Australia (#1124784, #1066770, #1057852, #1124907 to N.D.H; and #1140406 to F.S.F.G). F.S.F.G. was supported by a NHMRC Early Career Fellowship (1088703), a National Breast Cancer Foundation (NBCF) Fellowship (PF-15-008), a grant #1120725 awarded through the Priority-driven Collaborative Cancer Research Scheme and funded by Cure Cancer Australia with the assistance of Cancer Australia. N.D.H is a NHMRC CDF2 Fellow (1124788), a recipient of a Melanoma Research Grant from the Harry J Lloyd Charitable Trust, Melanoma Research Alliance Young Investigator Award, Tour De Cure research grant, equipment grant from The Ian Potter Foundation and a CLIP grant from Cancer Research Institute. This study was made possible through Victorian State Government Operational Infrastructure Support and Australian Government NHMRC Independent Research Institute Infrastructure Support scheme. **Author contributions:** A.I.W., C.C.O., C.H., D.S.H., J.R., I.S.S., J.C., L.D., M.D., M.A.D-E., R.D., R.H., S.H.Z., and F.S.F.G. designed, performed research and analyzed data. E.V. provided key reagents and scientific input into interpretation of the results. N.D.H and F.S.F.G. supervised work and wrote the paper. **Competing interests:** N.D.H. and J.R. are cofounders and shareholders in oNKo-Innate. E.V. is a cofounder and shareholder in Innate Pharma. The other authors have declared that no conflict of interest exists. **Material availability:** Prof. Stefan Karlsson for providing the TgfbRII flox mice; and Dr. Badia Kita / Paranta Biosciences Ltd for clinical grade Follistatin for *in vivo* assays.

---

**Table S1:**
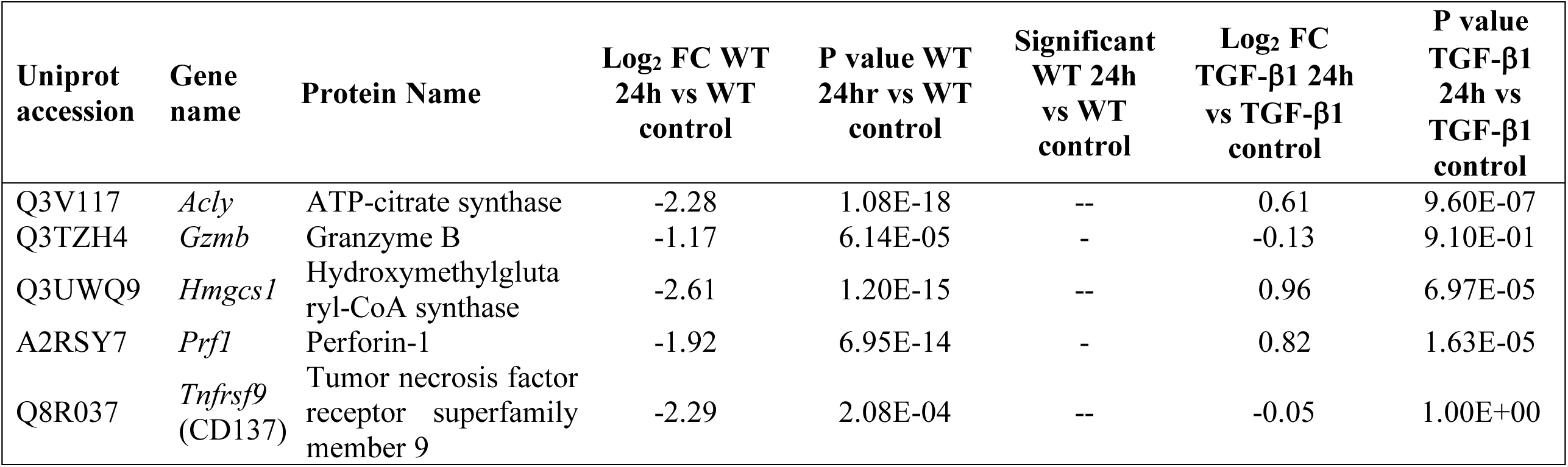
Summary of the Log_2_ FC and P values associated with the label-free quantitative proteomics experiments on WT and RII^FL^ samples comparing 24 h post rTGF-β1 stimulation vs unstimulated (control) cells. Significantly dysregulated proteins are indicated by “-” as a 2-fold or “--” 4-fold change.

**Figure S1:**
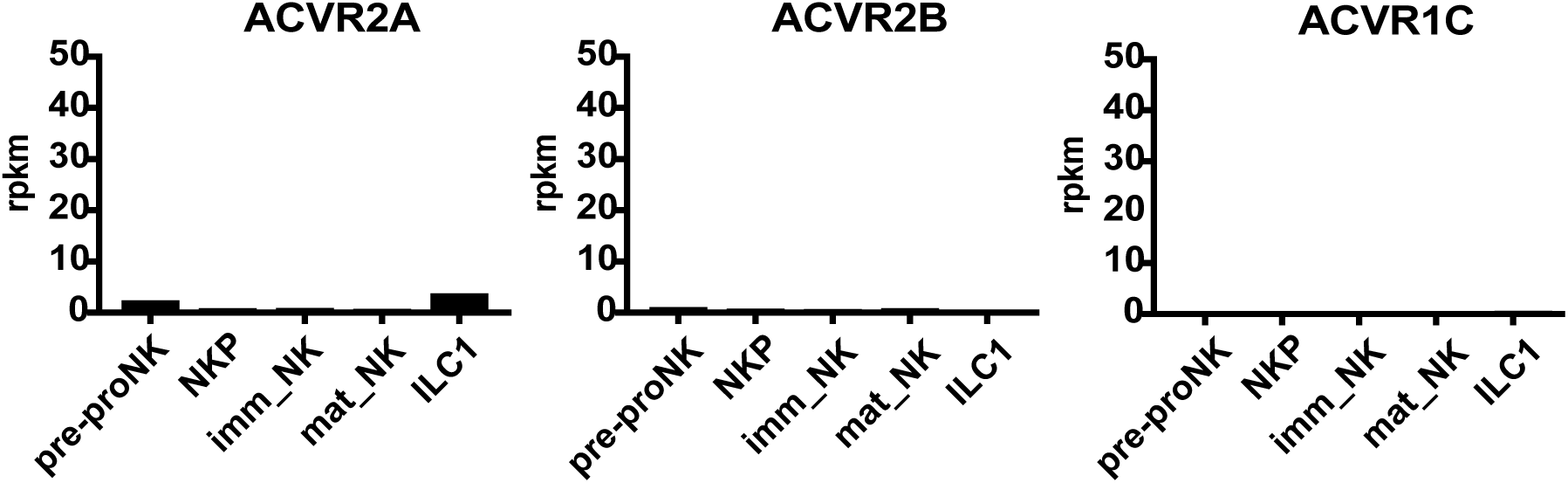
Alternative Activin receptors (ACVR2A, ACVR2B and ACVR1C) RNA expression. NK cell progenitors (NKP), immature (immat_NK) and mature (mat_NK) cells, and from the liver homogenate of a WT mice to sort innate lymphoid cells type 1 (ILC1: sorted by TRAIL^+^, Dx5^neg^, Lin^neg^), were subjected to RNAseq analysis (as previously shown (Delconte et al., 2016b; Holmes et al., 2014; Seillet et al., 2014)). Graphs show the reads per kilobase per million (RPKM) for each gene mapped to the indicated cell subsets.

**Figure S2:**
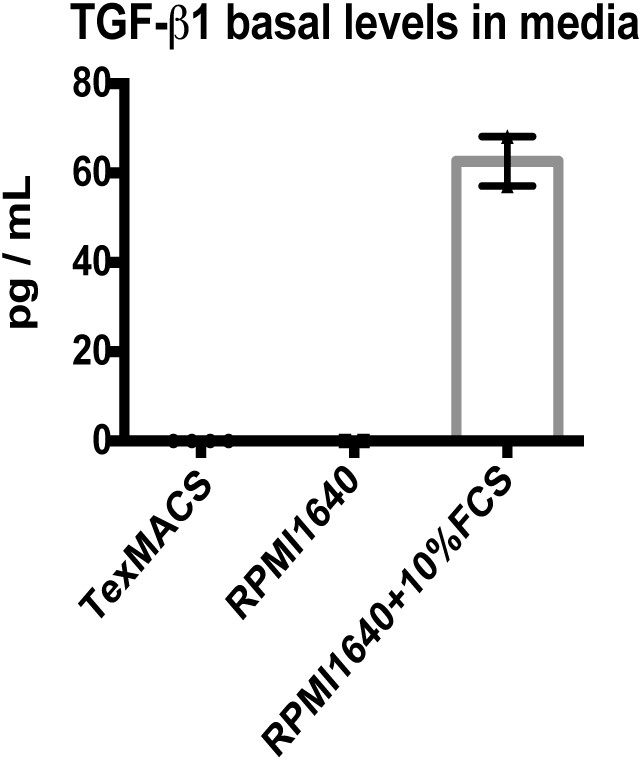
TGF-β1 levels in TexMACS animal free media, conventional RPMI 1646 and RPMI 1640 +10% fetal calf serum (FCS).

**Figure S3:**
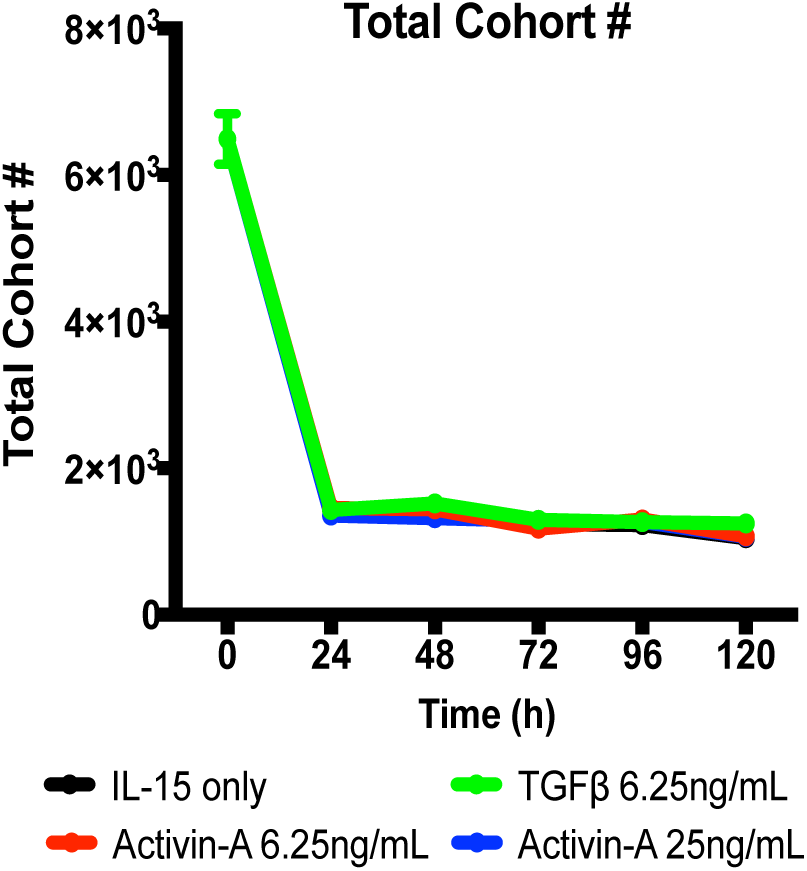
Total NK cell cohort number after Activin-A or TGF-b treatment. Freshly-isolated mouse splenic NK cells were labelled with a division-tracking dye and cultured in the presence of 50 ng/mL rIL-15 in addition to rTGF-β1 or rActivin-A as indicated. To measure of cell death or loss at each time point, the total cohort number (an estimation of the number of starting cells whose progeny represent the population at that time point) was calculated as described (Marchingo et al., 2014). Results shown are for NK cells isolated from one mouse but representative of three independent experiments. Error bars represent the S.D. of 3 technical replicates.

**Figure S4:**
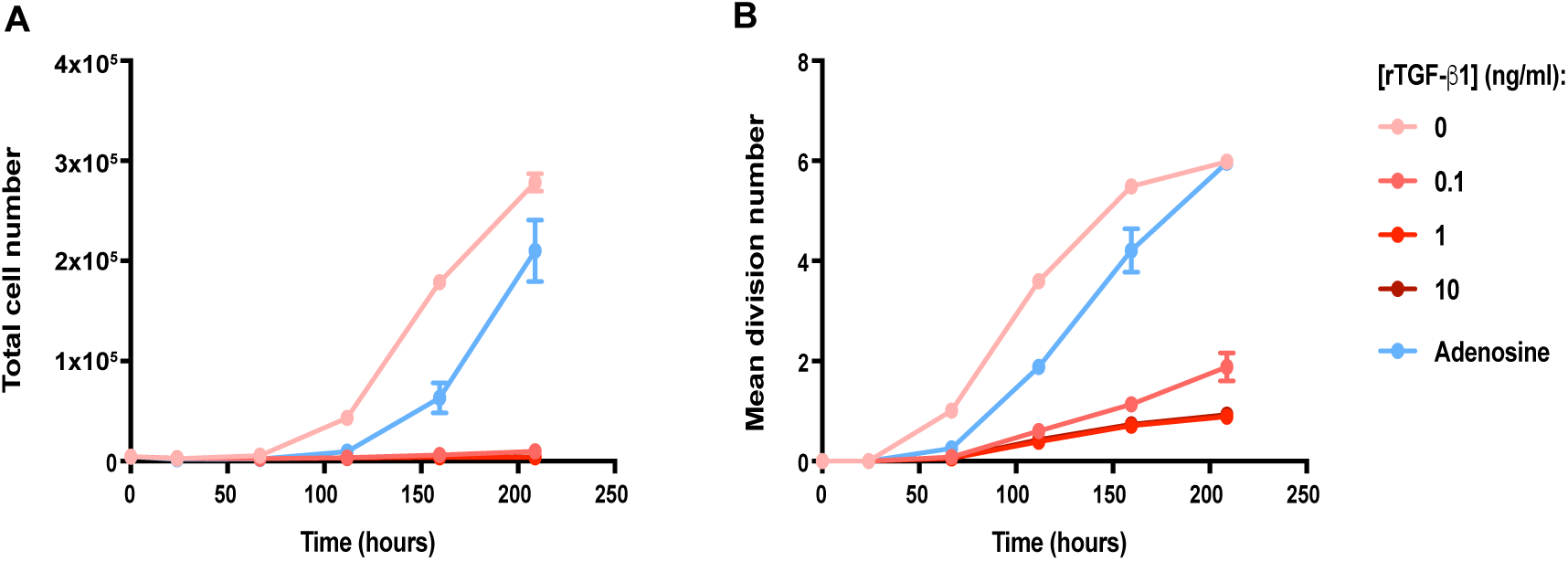
Comparison of adenosine and TGF-b treatments on NK cell proliferation. Splenic WT NK cells (live Lin^-^NK1.1^+^NKp46^+^CD49a^neg^CD49b^+^) sorted from spleens of WT mice were labelled with CFSE, and 8 x 10^3^ labelled cells were seeded into 96-well round bottom plates containing the indicated rTGF-β1 concentrations or adenosine (NECA) at 1 µM. Cells were analyzed by flow cytometry at the indicated time points and NK cell total (A) and mean division (B) numbers were determined using the Precursor Cohort Based Method measuring lymphocyte proliferation, survival and differentiation using CFSE time-series data (Hawkins et al., 2007; Marchingo et al., 2014). Data is representative from 2 independent experiments.

**Figure S5:**
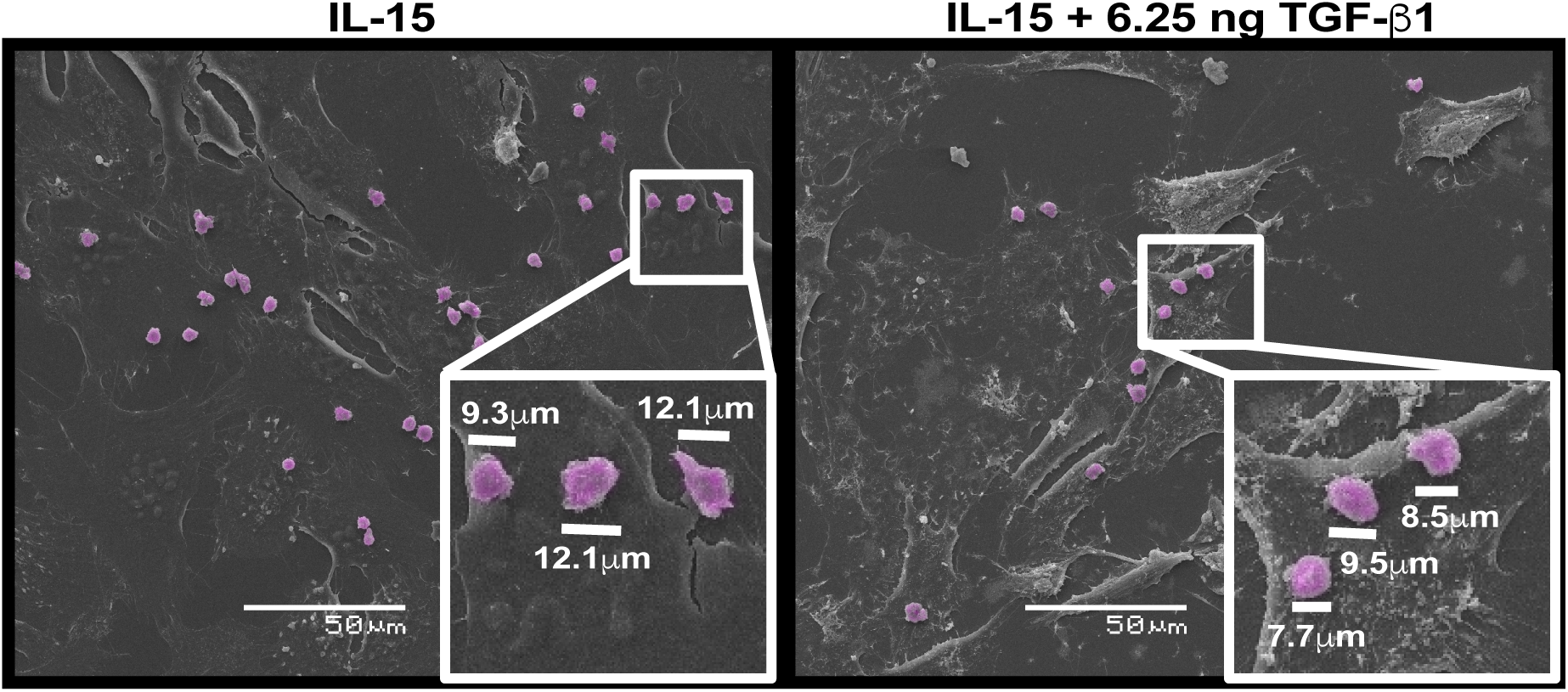
Representative scanning electron micrographs of melanoma cells adhered on glass coverslips and co-cultured with sorted NK cells overnight (16 h) in presence of 50 ng/mL rIL-15 with or without TGF-β1 (6.25 ng/mL) in animal free media. NK cells are digitally colored and indicated in magenta, and scale bars indicate the cellular dimensions of 3 NK cells per group. Images are representative from 3 different biological replicates.

**Figure S6:**
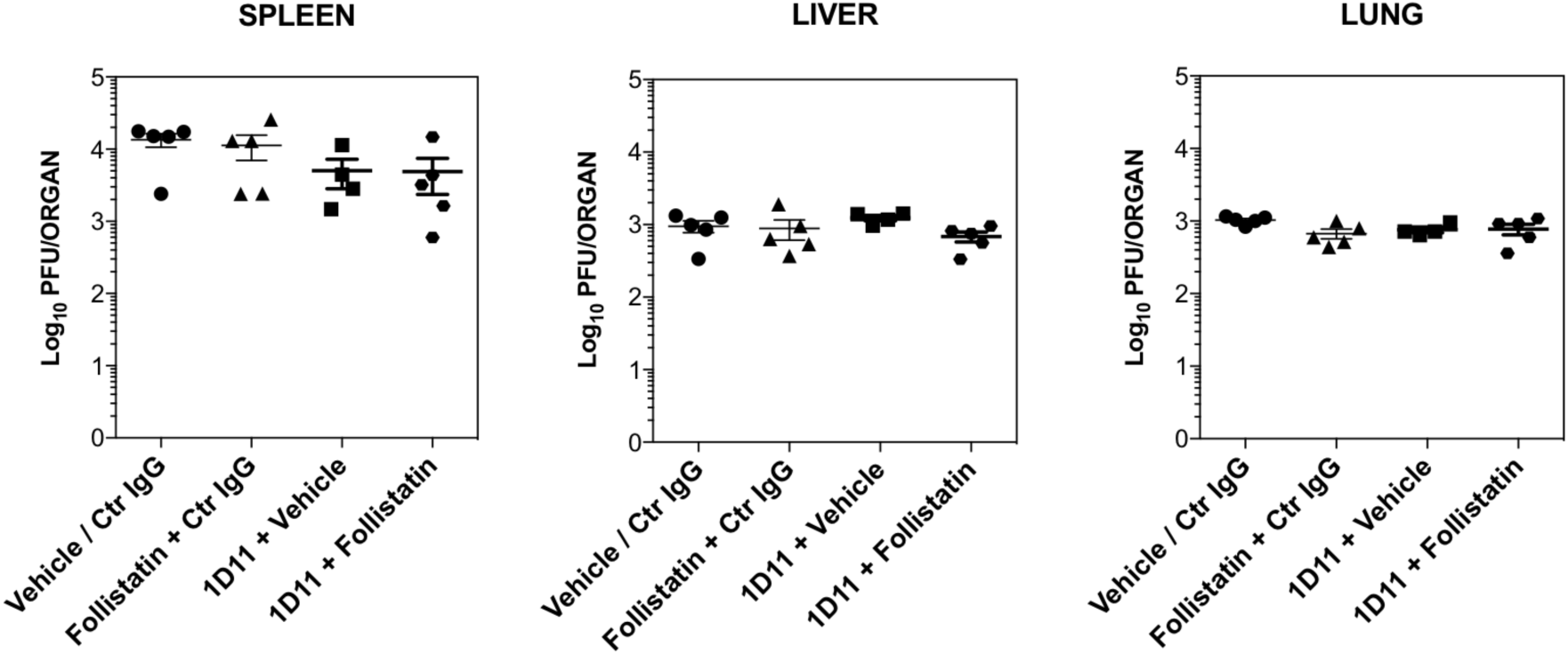
MCMV viral titters after anti-TGF-β1,2,3 or Follistatin treatment. Groups of 5 BALB/c mice were inoculated intraperitoneally (i.p.) with MCMV for 6 days. Mice were then i.p. treated with Follistatin (10 µg) and anti-TGF-β1,2.3 neutralizing antibody (250 µg) daily from d0 to endpoint. Organs (spleen, liver and lungs) were harvested for quantification of viral titter on day 6 post infection. Symbols in scatter plots represent the number of PFU per organ from individual mice (with mean and SEM shown by cross-bar and errors). Mann-Whitney test was used to compare differences between groups.

**Figure S7:**
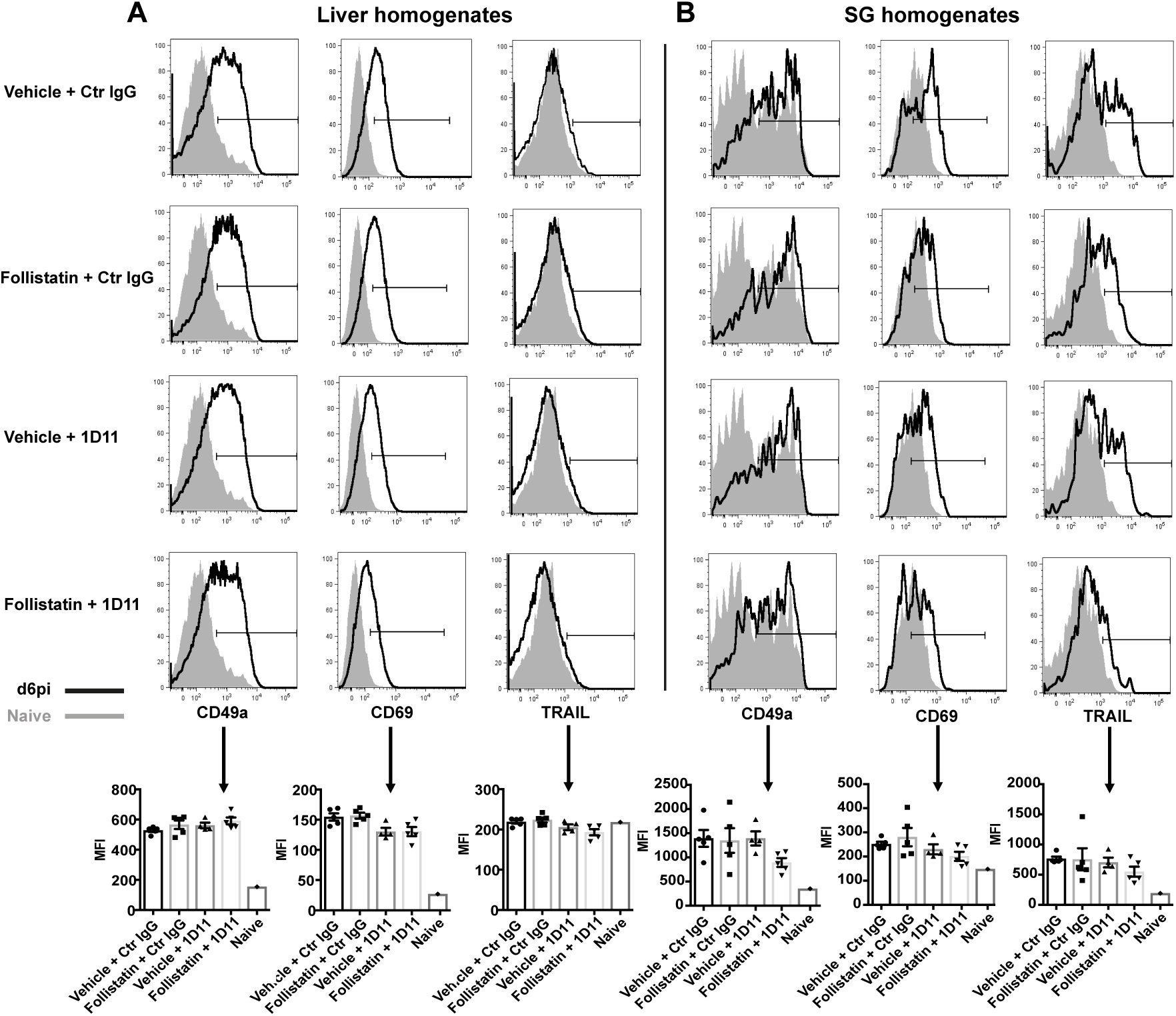
CD49a, CD69 and TRAIL expression at steady state or 6 days post MCMV infection in NKp46^+^ cells of liver (A) or salivary gland homogenates (B). Data is representative from 5 different biological replicates, where histograms are overlays of one naive control or one biological replicate at day 6 days post infection (d6pi) biological replicate. The expression of CD49a, CD69 and TRAIL for groups treated with Follistatin or neutralizing anti-TGF-β1,2,3 (1D11) antibody and the respectively controls (vehicle or ctr. IgG) was compared by quantitating the mean of fluorescence intensity (MFI). Symbols in scatter plots represent each individual mouse (with mean and SEM shown by cross-bar and errors). MannWhitney test was used to compare differences between groups.

**Figure S8:**
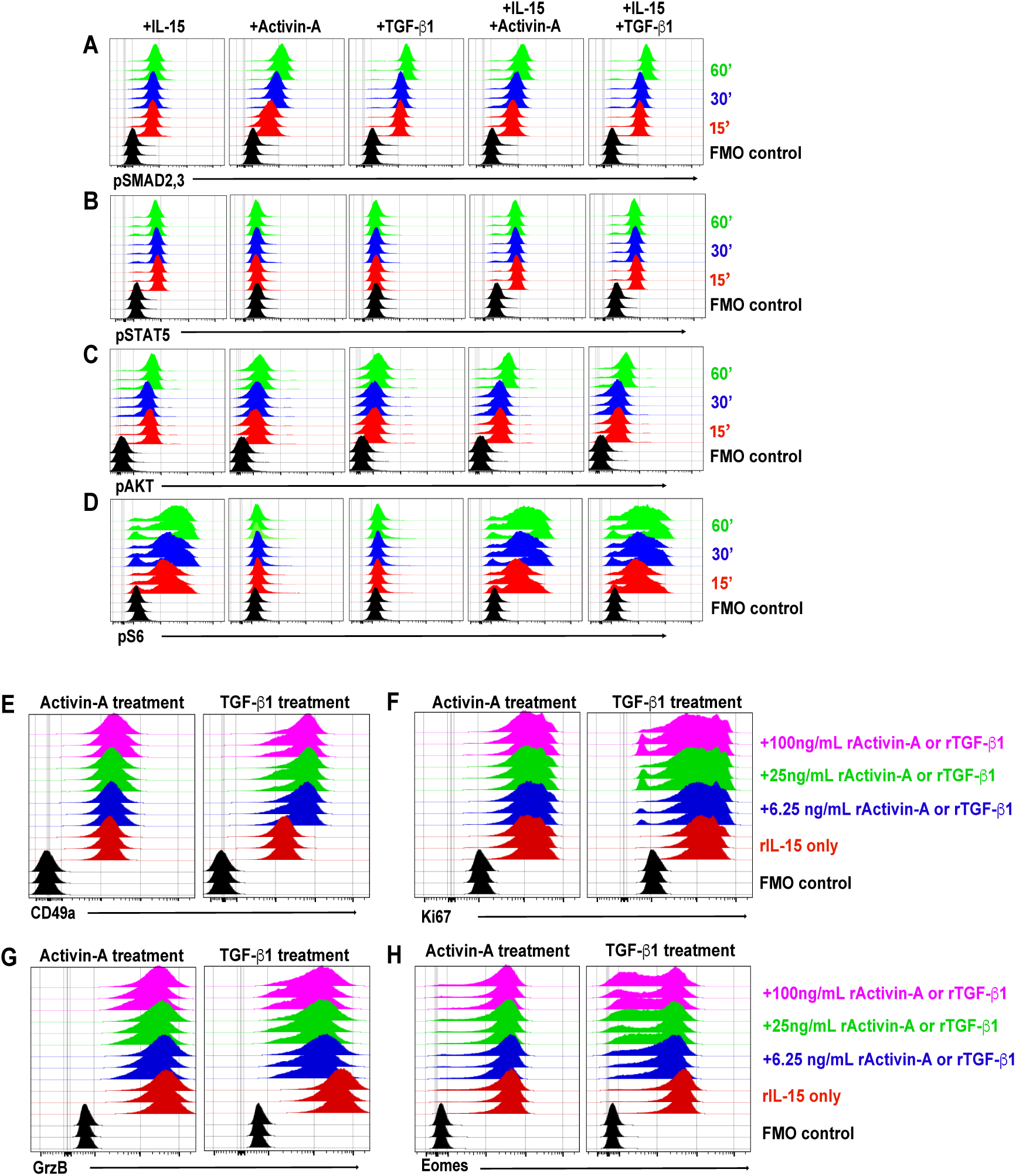
Conserved features of Activin signaling in hNK cells. NK cells were isolated from cord blood and primed in rIL-15 for 48 h. Cells were then washed, starved for 4 h in animal free media at 37°C, and stimulated in rIL-15 media with indicated concentrations of rActivin-A or rTGF-β1 for 1 h. pSMAD2,3 (A), pSTAT5 (B), pAKT (C) and pS6 (D) were analyzed by FACS. E-H: The same hNK samples, were separately cultured for 7 days in presence of the same culture conditions. At endpoint, cells were cell surface stained with anti-CD49a (E), and intracellularly stained with anti-Ki67 (F), anti-GrzB (G) and anti-Eomes (H) and analyzed by flow cytometry. Each overlay represents one representative independent biological replicate.

